# Caffeine-regulated molecular switches for functional control of CAR T cells *in vivo*

**DOI:** 10.1101/2025.06.13.659487

**Authors:** Elise Sylvander, Benjamin Salzer, Dominik Emminger, Hayeon Baik, Giulia D’Accardio, Markus Schäfer, Konstantina Mouratidis, Michelle C. Buri, Daniel Maresch, Anna Urbanetz, Jacqueline Seigner, Elisa Marchiori, Álvaro Muñoz-López, Fabian Engert, Boris Engels, Joerg Mittelstaet, Joris van der Veeken, Antonio Rosato, Johannes Zuber, Eva M. Putz, Charlotte U. Zajc, Manfred Lehner, Michael W. Traxlmayr

**Author notes:** These authors contributed equally to this work (Manfred Lehner, Michael W. Traxlmayr).

## Abstract

The limited controllability of CAR T cells in patients represents a key challenge of this highly potent immunotherapy. A molecular ON-switch, which can be regulated with a non-toxic and readily available small molecule drug, would represent a major advance towards controllable CAR T therapeutics. For that purpose, we engineered caffeine-responsive heterodimeric ON-switches (CaffSwitches) and demonstrate their high caffeine-dependency and virtually absent leakiness. When incorporating these CaffSwitches into CARs, the resulting CaffCARs were efficiently activated by caffeine concentrations achieved in human plasma after drinking one cup of coffee. Moreover, CaffCAR T cells also showed efficient tumor clearance in an *in vivo* mouse model, which was completely abolished in the absence of caffeine. This tight control was even observed with c-Jun overexpressing CaffCAR T cells, despite their massive expansion. Together, we anticipate that these novel CaffSwitches will be valuable tools for the development of safe and efficient next generation CAR T cells.

## Introduction

Despite their impressive clinical success^1–7^, currently approved chimeric antigen receptor (CAR) T cells still suffer from a number of limitations. For example, while the long-term proliferation and persistence of CAR T cells in the patient is beneficial for efficient tumor control^8–10^, their uncontrolled expansion becomes highly problematic in the case of severe toxicities. Importantly, these safety concerns are even increased with next generation CAR T cells that are additionally engineered to enhance potency and/or persistence, for example by overexpressing the AP-1 factor c-Jun^11^ or disruption of *TET2*^12^.

Approaches to overcome this lack of controllability after administration to the patient include the regulation of CAR function with dasatinib^13,14^ or with molecular switches, also known as chemically induced dimerization (CID) systems^15^, in which interaction of two proteins is induced with a small molecule drug. While several promising switches have been introduced, their application is limited by undesired pharmacological effects or toxicities of the small molecule drug^16–20^, potential immunogenicity of non-human protein components^21–24^ and/or lack of availability or high costs of the small molecule^17,19^. Thus, there is a need for clinically applicable molecular switches that can be regulated by safe and readily available small molecule drugs.

We hypothesized that such a switch system could be built based on a previously generated caffeine-specific single-domain antibody (nanobody termed anti-caffeine VHH-WT, acVHH-WT) from llamas^25^, which was found to homodimerize upon binding to caffeine^26–30^. This extensively studied small molecule has highly favorable pharmacokinetic properties, including efficient penetration of tissues such as the brain, a half-life of ∼5 h and lack of toxicity when consumed at reasonable concentrations^31–33^. Caffeine is also broadly available and highly cost-efficient, facilitating straightforward clinical translation. Finally, high-dose caffeine is even used to treat apnea of prematurity in infants^34,35^, further highlighting the safety of this drug.

However, the caffeine-dependent system described above comes with two critical limitations: First, the fact that it induces homodimerization limits its applicability, since most applications require conditional dimerization of two different subunits, i.e. heterodimerization. Second, considerable leakiness has been observed in the absence of caffeine^30^, which we also confirmed in our experiments described below.

Therefore, by using high-end protein engineering, we adapted this acVHH-based system to yield improved switches which show (i) defined heterodimerization in response to caffeine, (ii) virtually absent background activation and (iii) adapted caffeine sensitivity to respond to clinically relevant concentrations. Moreover, we demonstrate that these improved switches enable efficient control of CAR T cell activity both *in vitro* and in a mouse tumor model *in vivo*. Remarkably, even with hyperproliferating CAR T cells that co-express c-Jun, anti-tumor activity was completely abolished in the absence of caffeine, highlighting the strict caffeine-dependency of these novel engineered switches.

## Results

### Engineering caffeine-dependent heterodimeric switches without background activation

To create defined heterodimeric switches, two successive engineering steps were necessary: (i) abolishing homodimerization of the original acVHH-WT nanobody and (ii) engineering a second acVHH variant as a caffeine-dependent heterodimerization partner.

To eliminate homodimerization, we introduced charged amino acids into the hydrophobic homodimerization interface of acVHH-WT, yielding four different acVHH variants (Suppl. Fig. 1A). We chose acVHH-V106D based on its favorable biochemical properties (strictly monomeric behavior and lack of aggregation, Suppl. Fig. 1A) and due to the lack of homodimerization in the presence and absence of caffeine (Suppl. Fig. 1G).

Next, to engineer a caffeine-dependent heterodimerization partner for this acVHH-V106D variant, we adapted the interaction surface on the opposing nanobody by randomly mutating eight amino acid positions on a loop that is in close proximity to the V106D mutation (Fig. 1A). This library termed acVHH-8NNK was selected for binding to acVHH-V106D in the presence of caffeine by using the yeast surface display technology^36–39^ (Fig. 1B and Suppl. Fig. 1B). To ensure caffeine-dependency, multiple negative selections were performed by selecting for non-binding to acVHH-V106D in the absence of caffeine (Suppl. Fig. 1C and 1D). This engineering campaign ultimately yielded two acVHH variants termed acVHH-M1 and acVHH-M2, which expressed efficiently (Suppl. Fig. 1E) and showed high affinities to acVHH-V106D of 44 and 35 nM, respectively (Suppl. Fig. 1F). Importantly, the interaction of both variants with acVHH-V106D was strongly dependent on caffeine, with no detectable background activation (Fig. 1C), thus yielding two caffeine-responsive molecular switches (CaffSwitches). This is in contrast to acVHH-WT, which does show detectable caffeine-independent homodimerization (Suppl. Fig. 1H), as was also described in literature^30^. We further measured binding to acVHH-V106D in the presence of various caffeine metabolites and other structurally related endogenous molecules including adenine and guanine (Fig. 1C). While acVHH-M2 was triggered to some extent by all caffeine metabolites (theobromine, paraxanthine, theophylline), binding of acVHH-M1 was only induced by theophylline. Of note, contrary to theobromine, which is also found at high concentrations in chocolate^40,41^, the presence of theophylline is highly dependent on caffeine consumption. In summary, given its high dependency on caffeine with undetectable background activation and its non-responsiveness to theobromine found in chocolate, we chose acVHH-M1 for further experiments.

**Figure 1:**
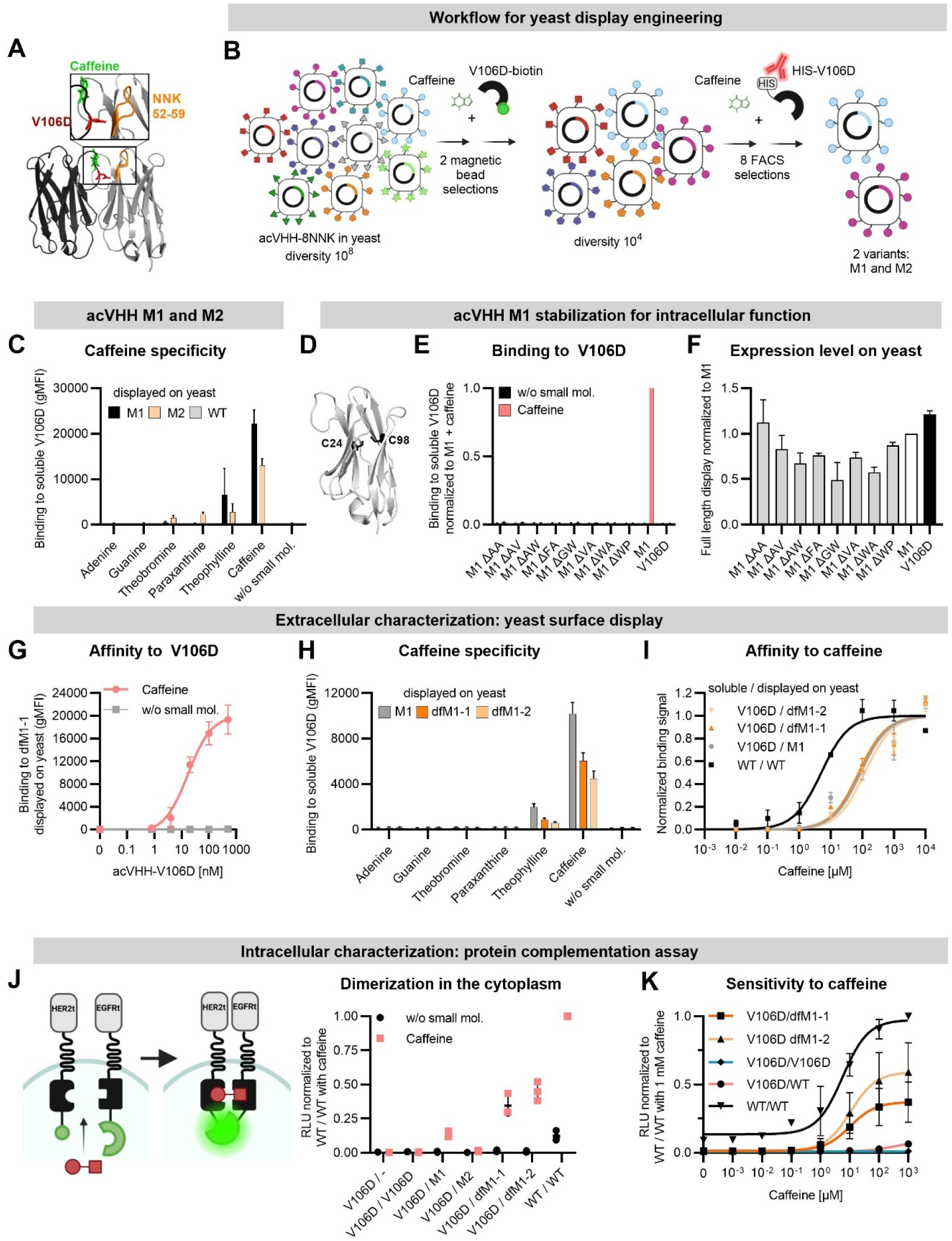
Engineering efficient and non-leaky CaffSwitches that are functional intracellularly. (A) Structure of the acVHH-WT dimer in complex with caffeine (PDB-ID 6QTL^27^). Positions that were mutated during engineering of the heterodimerizing acVHHs are highlighted. (B) Engineering process for the generation of caffeine regulated heterodimers. (C) Binding of three yeast-displayed acVHH variants to 500 nM soluble acVHH-V106D was assessed in the absence or presence of various small molecule compounds (10 µM) as indicated (mean ± SD, n=3). (D) Cartoon representation of acVHH-WT (PDB-ID 6QTL^27^) with the cysteines C24 and C98 highlighted in black. (E) Binding of acVHH-M1 variants containing deleted disulfide bonds to 500 nM soluble acVHH-V106D ± 10 µM caffeine. Mutated Cys positions are indicated with ΔXY, wherein X and Y indicate the mutations at positions C24 and C98, respectively. The geometric mean fluorescence intensity (gMFI) was normalized to acVHH-M1 with caffeine (mean ± SD, n=3). (F) Full-length display levels on yeast of disulfide-free acVHH-M1 variants shown in **(E)**. acVHH-M1 and -V106D were used as controls and the gMFI was normalized to acVHH-M1 (mean ± SD, n=3). (G) Binding of the yeast-displayed dfM1-1 to increasing concentrations of soluble acVHH-V106D was assessed ± 30 µM caffeine (mean ± SD, n=3). (H) Binding of three yeast-displayed acVHH variants to 500 nM soluble acVHH-V106D was assessed in the absence or presence of various compounds (10 µM) as indicated (mean ± SD, n=3). (I) Binding of yeast-displayed acVHH mutants to 400 nM soluble acVHH-V106D or -WT (as indicated) was measured in the presence of increasing concentrations of caffeine (mean ± SD, n=3). (J) Intracellular binding in the absence or presence of 30 µM caffeine was measured with the NanoBiT® protein complementation assay in Jurkat T cells and normalized to the acVHH-WT homodimer in the presence of caffeine (mean ± SD, n=3). RLU, relative light unit. (K) Intracellular binding of the acVHH variants in the presence of increasing concentrations of caffeine was measured with the NanoBiT® protein complementation assay in Jurkat T cells. The signal was normalized to the acVHH-WT homodimer in the presence of 1 mM caffeine (mean ± SD, n=4).

### Engineering disulfide-free CaffSwitches to enable intracellular functionality

Since engineering processes for improved protein functions are often associated with a loss in protein stability^42–44^, we speculated that mutating eight amino acid positions in acVHH-M1 slightly destabilized the protein and therefore increased dependency on its internal disulfide bond (Fig. 1D). As disulfides typically fail to form in the reducing environment of the cytoplasm, this would lead to misfolding and impaired activity when being expressed intracellularly^45–47^. Indeed, while the acVHH-M1-based CaffSwitch was functional under oxidizing conditions (Fig. 1C and Suppl. Fig. 1F), it lost much of its activity in the cytoplasm of Jurkat T cells (Suppl. Fig. 1I). Therefore, we mutated these Cys positions to amino acid pairs previously shown to be acceptable substitutions for disulfide bonds^48–50^. However, this ablated the function of the CaffSwitches (Fig. 1E), reaffirming the strong disulfide bond dependency of this nanobody.

To be able to apply our CaffSwitches also intracellularly, we engineered disulfide-free variants of acVHH-M1 by exchanging the Cys residues to amino acid combinations that yielded the highest expression levels on the yeast surface (AA, AV, VA or WP; Fig. 1F). This oligoclonal pool of disulfide-free nanobodies was further randomly mutated by error-prone PCR (epPCR), followed by multiple rounds of selection for efficient switch function (Suppl. Fig. 2A and 2B).

This engineering strategy finally yielded two mutants termed “disulfide-free M1-1” (dfM1-1) and dfM1-2, both of which were expressed efficiently (Suppl. Fig. 2C) and showed high affinities to acVHH-V106D of ∼20 nM (Fig. 1G and Suppl. Fig. 2D and 2G). Most importantly, their interaction with acVHH-V106D remained strongly caffeine-dependent, as demonstrated by the total lack of binding in the absence of caffeine (Fig. 1G and 1H). The improved CaffSwitches were selectively activated by caffeine and its metabolite theophylline but remained unresponsive to theobromine or to structurally related molecules such as adenine or guanine (Fig. 1H).

Another critical parameter of a small molecule-regulated switch is the concentration of the drug required for activation. For caffeine, high sensitivity is undesired, since low background levels of caffeine would already trigger the switches, even when trying to avoid intake of caffeine-containing food and beverages. Therefore, we used relatively high concentrations of caffeine (10 µM) during the engineering process to reduce the sensitivity to caffeine. Indeed, when compared to acVHH-WT, the *EC*_50_ values of the newly engineered CaffSwitches (based on acVHH-M1, dfM1-1 and dfM1-2) were elevated by ∼20-fold (Fig. 1I and Suppl. Fig. 2G).

To test whether the removal of the disulfide bond enables intracellular activity, we assessed switch function in the reducing environment of the cytoplasm of Jurkat T cells. Indeed, the disulfide-free CaffSwitches based on dfM1-1 or dfM1-2 exhibited enhanced caffeine-dependent dimerization compared with the disulfide-containing acVHH-M1 (Fig. 1J). In agreement with the yeast display experiments (Fig. 1G, 1H and 1I), background binding in the absence of caffeine was not detected in the cytoplasm of Jurkat T cells either (Fig. 1J, 1K and Suppl. Fig. 2F), demonstrating the tight control of these switches. Of note, this is in stark contrast to the original acVHH-WT homodimeric switch, which – in agreement with a previous study^30^ – showed considerable caffeine-independent dimerization (Fig. 1J and 1K). Moreover, these intracellular assays showed that responsiveness to chocolate-contained theobromine was eliminated during the engineering process (Suppl. Fig. 2E) and again confirmed the reduced sensitivity to caffeine (Fig. 1K and Suppl. Fig. 2F and 2G).

To sum up, we successfully engineered heterodimeric CaffSwitches which (i) exhibit no detectable leakiness, (ii) display reduced caffeine sensitivity, thus minimizing unintended activation by trace caffeine levels in humans, (iii) are unresponsive to theobromine present in high quantities in chocolate, and (iv) retain functionality in the intracellular environment.

### Caffeine-mediated regulation of CAR T cell activity

Based on these promising biochemical experiments, we next incorporated the dfM1-1- and dfM1-2-based CaffSwitches into split CD19-BBζ CARs (Fig. 2A). Different split CAR architectures containing monomeric or dimeric versions of either chain were tested in primary human T cells. All constructs showed caffeine dependency, in particular those with dimeric chain II (Fig. 2A). Remarkably, these split CARs based on dimeric chain II showed more pronounced switch behavior compared to a benchmark FKBP/FRB-based CAR^17^ and reached target cell lysis levels comparable to that of the conventional CD19-CAR (Fig. 2A). Finally, we also confirmed that target cell lysis was dependent on both caffeine and antigen recognition since no lysis of CD19-negative cells was observed (Fig. 2A).

**Figure 2:**
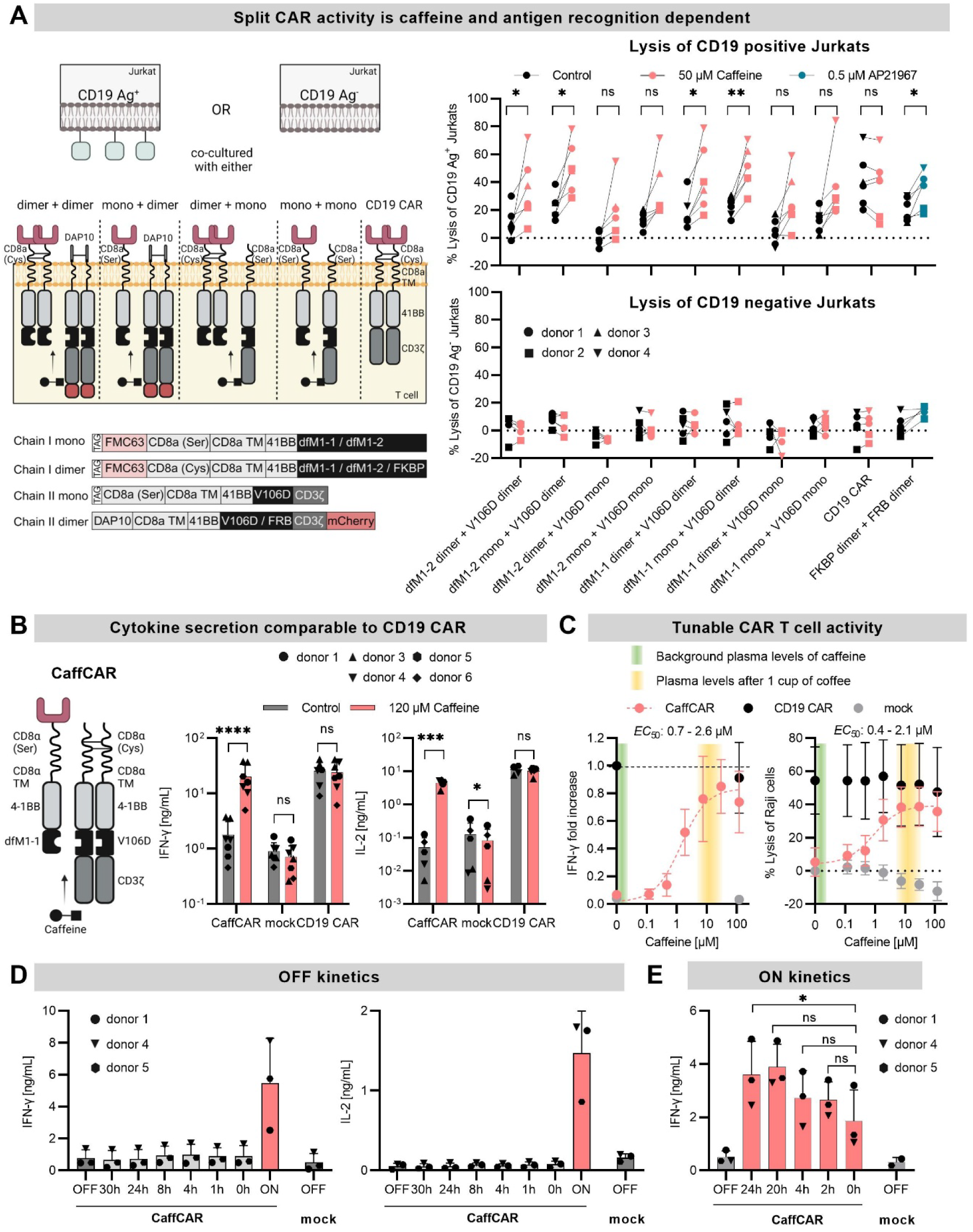
CaffSwitches enable efficient regulation of CAR function. **(A)** On the left, schematics of the different split CAR architectures are shown. For the dimeric chain I (chain I dimer), the original CD8α hinge containing 2 cysteines was used. Mutating these to serines resulted in monomeric chain I (chain I mono). The monomeric chain II (chain II mono) involves an extracellular tag fused to a CD8α hinge with cysteine to serine mutations. To dimerize chain II (chain II dimer), the CD8α hinge was exchanged with DAP10. The mCherry was fused C-terminally after the CD3ζ domain. The lysis of Jurkat T cells expressing or not expressing the CD19 antigen is depicted on the right. Expression of CAR chains in primary human T cells and of the CD19 antigen in Jurkat cells was achieved via mRNA electroporation. Co-cultures were set up with an E:T ratio of 2:1 for 4h (n = 6, 4 different T cell donors) and caffeine or the rapalog (AP21967) was added as indicated. **(B)** CAR T cells expressing the final CaffCAR (schematic on the left) were co-incubated with Raji cells (E:T of 2:1, 24 h), followed by analysis of IFN-γ (n = 7, 5 different T cell donors) and IL-2 (n = 5, 5 T cell donors). The values were also used in the titrations of Figure 2C for IFN-γ and Suppl. Figure 3C for IL-2. **(C)** Specific lysis of Raji cells (right) and IFN-γ secretion (left) were measured as a function of caffeine concentration (E:T of 2:1, 24 h). Lysis was normalized to mock T cells without caffeine and IFN-γ secretion was normalized to that of the CD19 CAR without caffeine (n = 7, 5 different T cell donors). **(D)** CaffCAR cells were cultivated with 30 µM caffeine (48 h before the co-culture) and subsequently washed to remove the drug at the indicated time points before setting up a co-culture with Raji cells without caffeine (E:T of 2:1, 24 h). The time points represent the time period spent in the absence of caffeine before co-culture. CaffCAR cells that were always in the presence of caffeine were washed once and caffeine was added (ON) or not added (0 h) in the co-culture. CaffCAR cells that were never cultured with caffeine (OFF) and mock T cells were used as negative controls. After 24 h, IFN-γ (left) and IL-2 (right) secretion were measured in the supernatants (n = 3, 3 different T cell donors). **(E)** 30 µM caffeine was added into CaffCAR T cell cultures at the indicated time points prior to the co-culture with Raji target cells. At t = 0 h a 4 h co-culture with Raji cells was set up (E:T of 2:1) with caffeine. CaffCAR cells that were never cultured with caffeine (OFF) and mock T cells were used as negative controls. After 4 h IFN-γ secretion was measured in the supernatants (n = 3, 3 different T cell donors). * < 0.05; ** < 0.01; *** < 0.001, **** < 0.0001. Multiple paired t-test (lysis) or ratio t-test (cytokine) with Holm Sidak correction.

Based on these observations, we chose a dfM1-1 split CAR design with monomeric chain I and dimeric chain II and further optimized it for clinical translation by excluding the potentially immunogenic mCherry marker. Furthermore, we replaced the extracellular domain of DAP10 with CD8α to reduce vector payload and confirmed that this replacement did not have an impact on split CAR function (Suppl. Fig. 3A). This final split CAR construct (Fig. 2B) termed “CaffCAR” showed efficient expression on primary human T cells (Suppl. Fig. 3B).

When tested in co-culture with CD19^+^ Raji lymphoma cells, CaffCAR T cells were potently activated in the presence of caffeine, yielding IFN-γ and IL-2 secretion levels comparable to those of conventional CD19 CAR T cells (Fig. 2B). Importantly, in the absence of caffeine, cytokine secretion was similar to mock T cells, again demonstrating the tight control of the engineered CaffSwitch. Caffeine dose-response experiments demonstrated that caffeine concentrations in the range of ∼5-15 µM are required for efficient cytokine secretion and cytotoxicity of CaffCAR T cells (Fig. 2C and Suppl. Fig. 3C). Of note, this corresponds to the caffeine concentration that is achieved in the plasma after drinking one cup of coffee^51,52^. Finally, kinetics experiments demonstrated that CaffCAR T cells respond quickly to caffeine withdrawal (Fig. 2D) and caffeine administration (Fig. 2E).

To test the versatility of the CaffSwitch in CAR T cells, we aimed to assess its switch function in CARs with different antigen specificities and costimulatory domains. For this purpose, we integrated the CaffSwitch into prostate specific membrane antigen (PSMA)-directed CARs containing either a 4-1BB-CD3ζ (BBζ) or CD28-CD3ζ (28ζ) backbone (Fig. 3A). When co-cultured with PC3 target cells engineered to express PSMA, as well as with their PSMA-negative counterparts (Fig. 3B), we demonstrated that cytotoxicity and cytokine secretion were highly caffeine- and antigen-dependent with both CAR backbones (Fig. 3C). Moreover, caffeine-dependent CaffPSMA-CAR function was also observed with LNCaP prostate cancer cells endogenously expressing the PSMA antigen (Fig. 3D).

**Figure 3:**
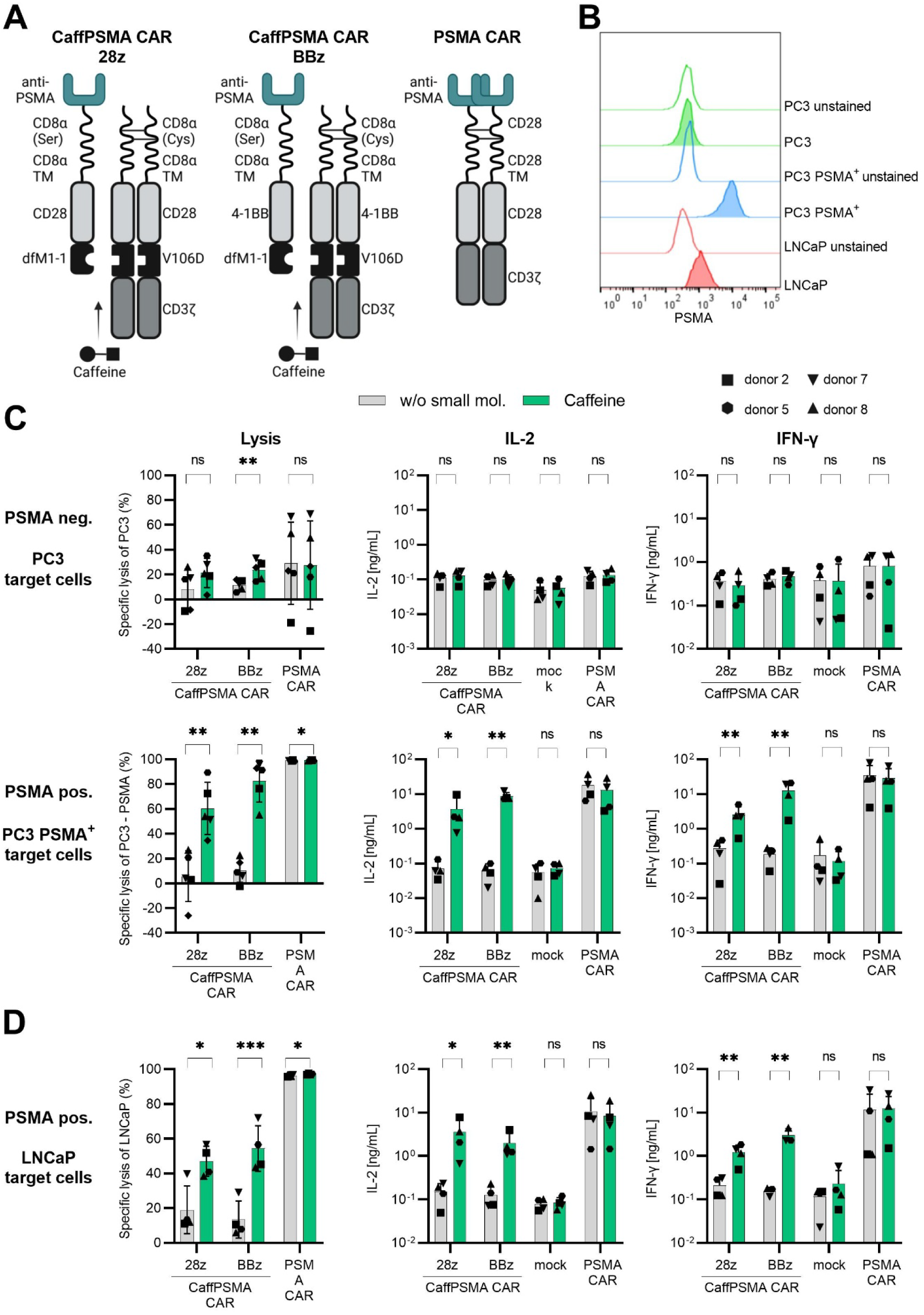
Efficient regulation of PSMA-directed CaffCARs based on different CAR backbones. **(A)** Schematic representation of the PSMA CaffCARs and a conventional control PSMA CAR. **(B)** PSMA expression levels on the different cell lines used in co-culture assays. **(C-D)** Specific lysis (left, n = 5), IL-2 secretion (middle, n = 4) and IFN-γ secretion (right, n = 4) after co-culture (E:T of 2:1, 24 h) with **(C)** PC3 cells engineered to express PSMA or non-engineered PC3 cells and **(D)** LNCaP cells endogenously expressing PSMA. Different symbols are used to represent the individual T cell donors. * < 0.05; ** < 0.01; *** < 0.001. Multiple paired t-test (lysis) or ratio t-test (cytokine) with Holm Sidak correction.

Together, we demonstrated that CaffCARs (i) reach activation levels comparable to those of conventional CARs, (ii) show tight control in the absence of caffeine, (iii) can be rapidly switched on and off, (iv) are adaptable to different antigen specificities and CAR backbones and (v) respond efficiently to caffeine plasma levels achievable with a single cup of coffee.

### Transient deactivation of CAR signaling by caffeine withdrawal limits T cell exhaustion

It has been shown previously that transient rest by either CAR degradation or pharmacologic inhibition with dasatinib reverses T cell exhaustion and restores effector functions^13,14^. To investigate whether exhaustion can also be prevented using CaffCARs, we conducted long-term co-cultures of CD19-BBζ CaffCAR T cells with Raji target cells, alternating between ON- and OFF-states (4–5-day intervals) by administering or withdrawing caffeine (Fig. 4A). Remarkably, despite being active only half the time, these transiently paused CaffCAR T cells (“CaffCAR pause”) exhibited enhanced expansion compared with continuously activated counterparts (“CaffCAR ON”) (Fig. 4B), as well as reduced expression of exhaustion-associated markers (PD-1, LAG-3, TIM-3, CD39; Fig. 4D and Suppl. Fig. 4A). This improved activation in the “CaffCAR pause” condition also translated to a reduced percentage of tumor cells in these co-cultures (Fig. 4C and 4F). At the end of the experiment, both co-cultures were split and incubated with or without caffeine for 5 days. Notably, adding caffeine to the “CaffCAR pause” culture elevated the expression of these markers to levels comparable with those of “CaffCAR ON +caffeine”, confirming efficient reactivation of CaffCAR T cells after transient rest. Likewise, even after 32 days of continuous activation (“CaffCAR ON”), a 5-day rest period (“CaffCAR ON -caffeine”) was sufficient to reduce exhaustion marker expression to levels similar to “CaffCAR pause -caffeine” (Fig. 4E and Suppl. Fig. 4B). Together, these data suggest that ON/OFF-cycles of split CAR activation limit CAR T cell exhaustion, thereby promoting CAR T cell expansion and tumor control.

**Figure 4:**
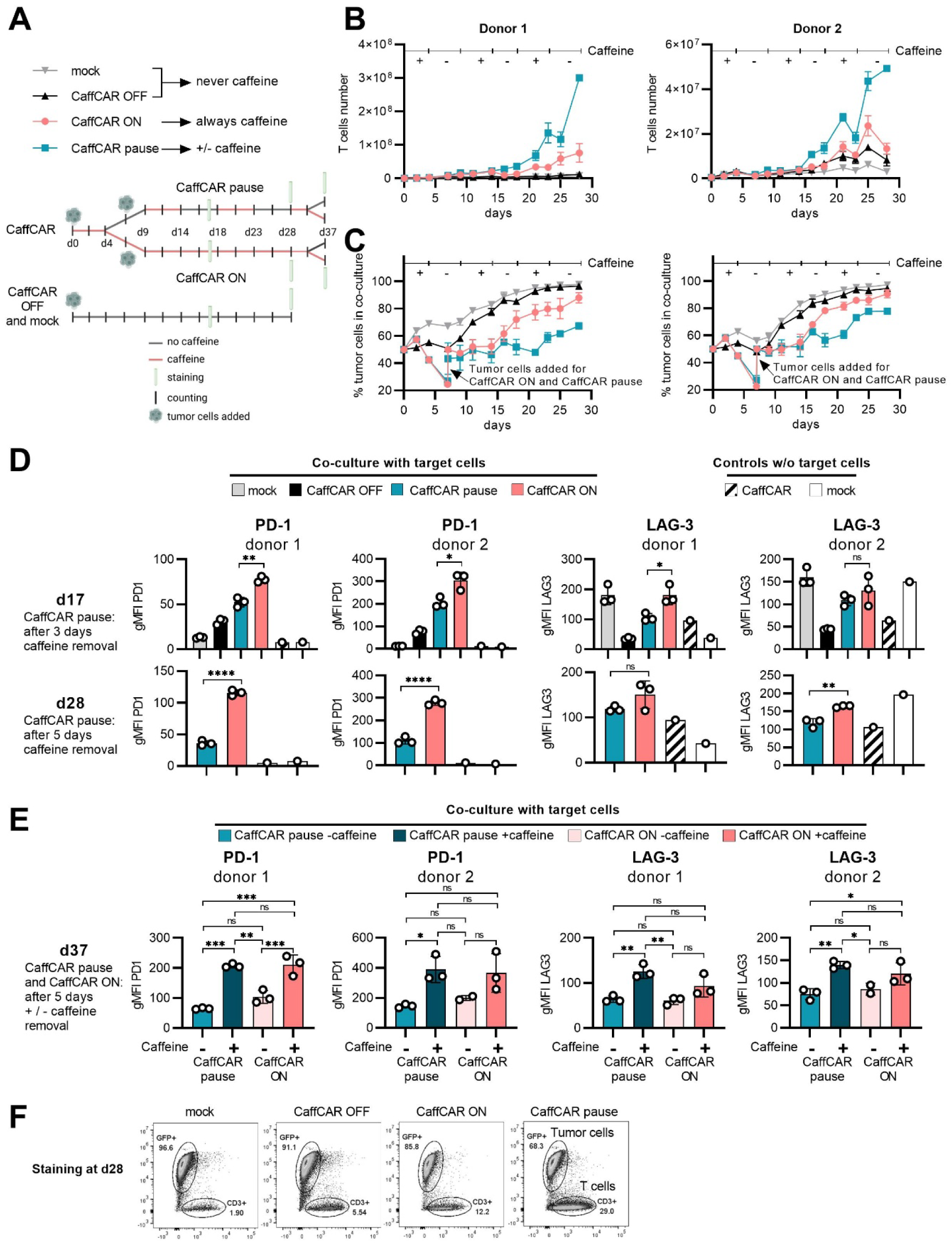
Pausing caffeine administration limits T cell exhaustion. **(A)** Schematic of the long-term co-culture experiment. **(B-C)** T cell expansion **(B)** and percentage of Raji cells in the co-cultures **(C)**. The presence or absence of caffeine in the co-cultures of the CaffCAR pause condition is indicated on the top. Technical triplicates for two independent T cell donors are shown. **(D-E)** Expression of PD-1 and LAG-3 on d17 and d28 **(D)** and d37 **(E).** Technical triplicates for two independent T cell donors are shown. On d28, mock and “CaffCAR OFF” T cells are not depicted due to insufficient T cell numbers in the co-cultures. **(F)** Representative flow cytometric measurements at the end of the co-culture. * < 0.05; ** < 0.01; *** < 0.001. Unpaired t-test with Welsh’s and Holm Sidak correction (D), and 1-way Anova with Turkey’s test for d37 (E).

### Efficient functional control of CaffCAR T cells *in vivo*

Next, we assessed the function of CD19-BBζ CaffCAR T cells in a Raji lymphoma mouse model *in vivo* (Suppl. Fig. 5A). Administration of caffeine via drinking water significantly increased CaffCAR T cell expansion and tumor control (Suppl. Fig. 5B). However, we only observed long-term survival in 2/9 mice in the “CaffCAR+caffeine” group, indicating limited anti-tumor potency (Suppl. Fig. 5C). We hypothesized that the short half-life of caffeine in mice (0.5 h vs. 5 h in humans^31,53^), together with irregular drinking habits, might lead to periods of insufficient caffeine levels and thus inactivity of CaffCAR T cells. Indeed, plasma caffeine concentrations were below the limit of quantification (5 nM) in 4/9 mice at the day of sacrifice (Fig. 5A), approximately 1000-fold below the level required for effective CaffCAR activation (Fig. 2C).

**Figure 5:**
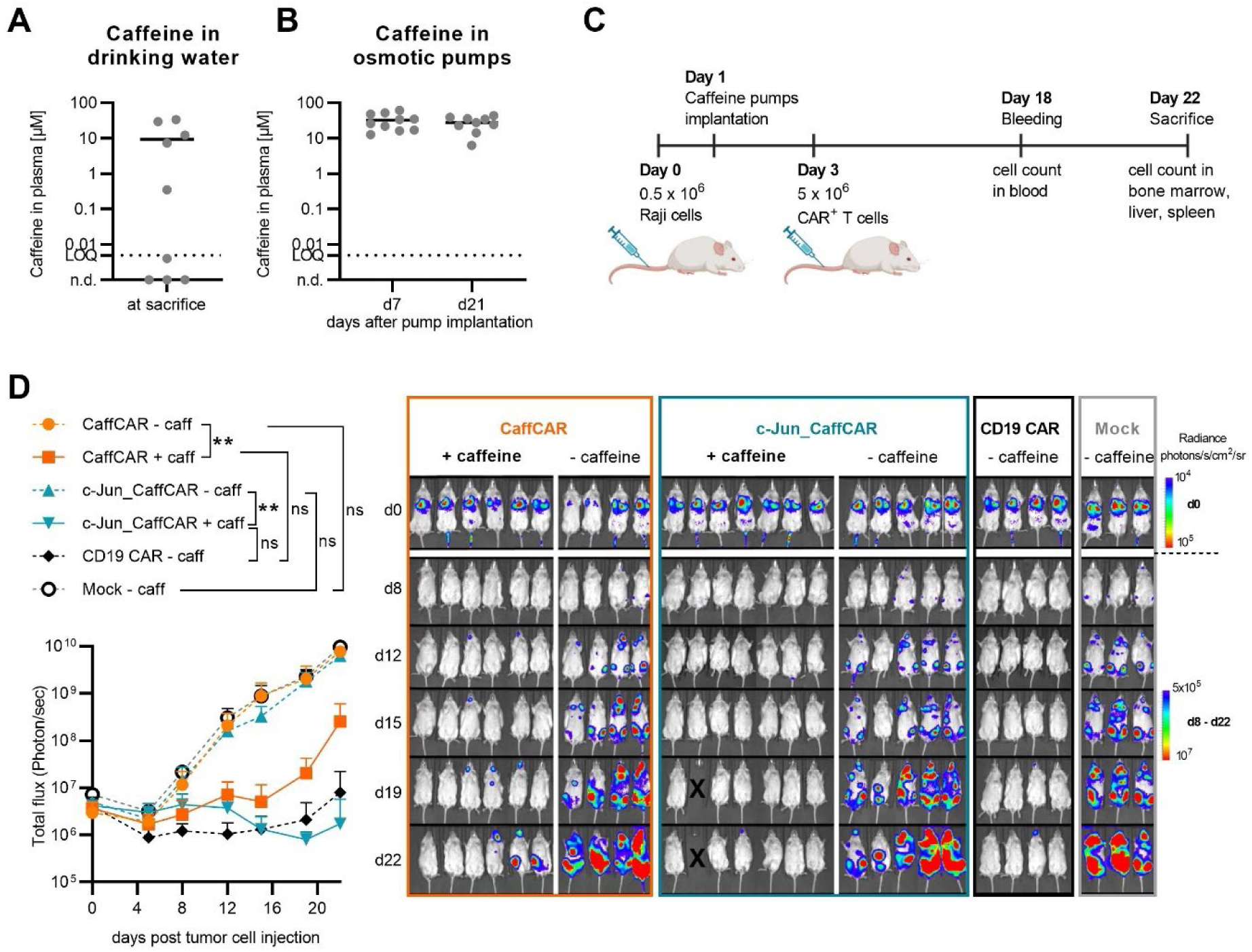
Caffeine tightly regulates CaffCAR function *in vivo*. **(A-B)** Mass spectrometric analysis of caffeine concentration in the plasma of mice who received caffeine via **(A)** the drinking water (n = 9) or **(B)** osmotic pumps, 7 and 21 days post pump implantation (n = 10). Limit of quantification (LOQ): 5 nM; non-detectable (n.d.). **(C)** Schematic of the *in vivo* experiment with osmotic pumps. **(D)** Tumor load as determined by BLI of the *in vivo* experiment with osmotic pumps, with quantified total flux on the left and pictures on the right. In the c-Jun_CaffCAR group, one mouse (tumor free according to BLI signal) needed to be sacrificed due to tumor-unrelated complications (reopening of the wound, where the pump was implanted). * < 0.05; ** < 0.01. 2-way Anova with Turkey’s test.

To achieve more stable and elevated caffeine levels, we switched from administration via drinking water to osmotic pumps, which provide continuous caffeine release. Consistently high plasma caffeine concentrations were confirmed in all mice in a pilot experiment (Fig. 5B).

Based on these findings, pumps were implanted into lymphoma-bearing mice prior to CAR T treatment. To assess whether CaffCARs also enable control of boosted CAR T cells engineered for enhanced proliferation and exhaustion resistance, we additionally tested CaffCAR T cells overexpressing c-Jun (Fig. 5C, 5D; Suppl. Fig. 5D). Strikingly, tumor control by CaffCAR T cells — both with and without c-Jun overexpression — was strongly dependent on caffeine administration (Fig. 5D). While CaffCAR T cells reduced tumor burden effectively, c-Jun overexpression further enhanced their potency, achieving anti-tumor activity comparable to or exceeding that of conventional CD19 CAR T cells. In contrast, without caffeine, tumor growth in both CaffCAR groups (with or without c-Jun) resembled that of the mock T cell group, confirming the absence of leakiness (Fig. 5D). These findings demonstrate that CaffSwitches enable tight *in vivo* control of CAR T cells, including those potentiated via c-Jun overexpression.

### c-Jun drives CAR-independent T cell hyperproliferation

Unexpectedly, solid white nodules developed in the livers of most mice treated with c-Jun_CaffCAR T cells, which was not observed in any of the other mice (CaffCAR, CD19 CAR or mock T treated), suggesting CAR T cell hyperproliferation (Fig. 6A). Importantly, nodule formation occurred regardless of caffeine administration, indicating that CAR signaling via CD3ζ was not required.

**Figure 6:**
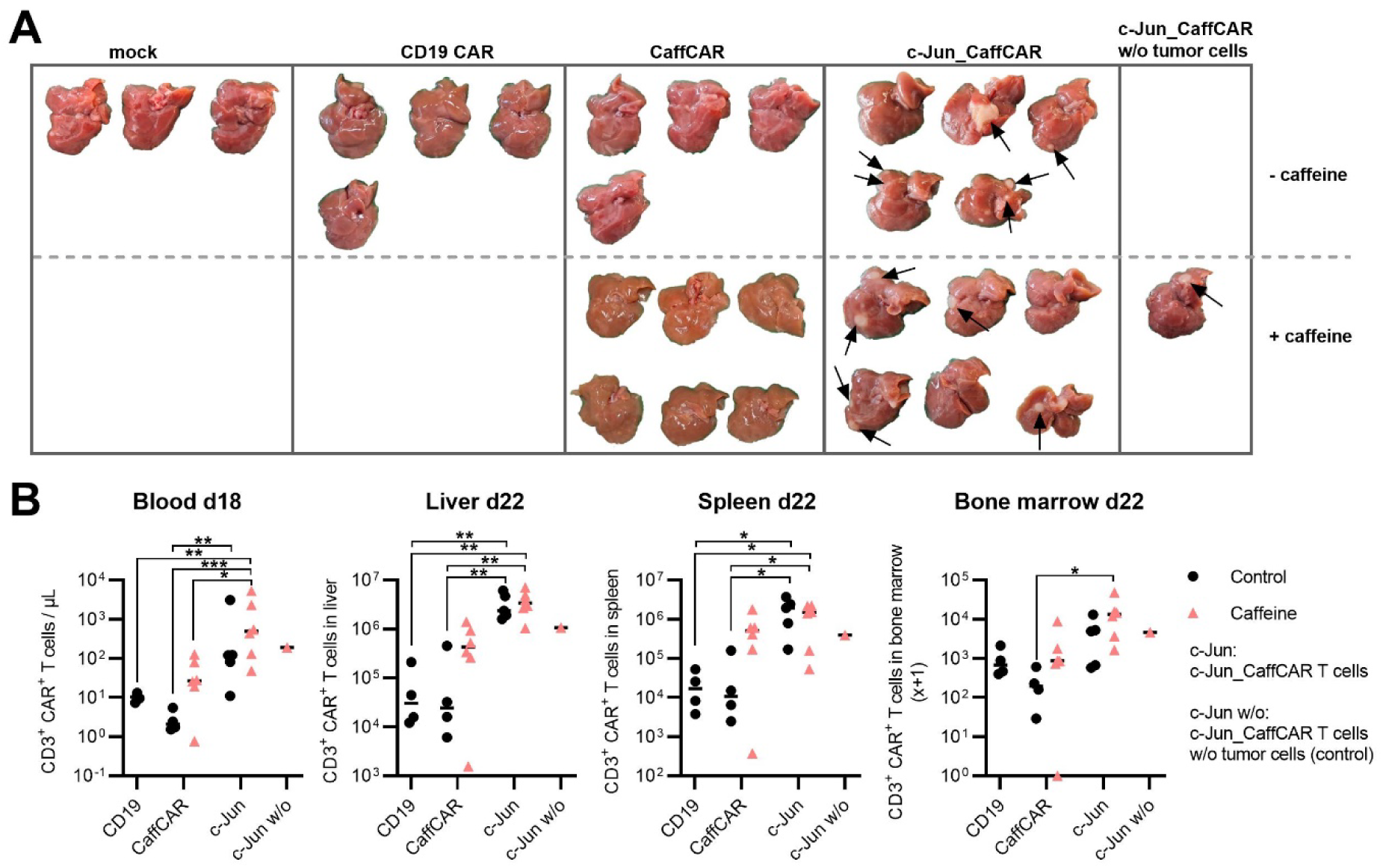
c-Jun drives CAR T cell hyperproliferation. (A) Livers of all mice at sacrifice. The arrows point towards solid white nodules. (B) Quantification of CAR^+^ T cells in the blood on day 18 (d18) and in the liver, spleen and bone marrow on d22. * < 0.05; ** < 0.01. 1-way Anova with Turkey’s test.

Strikingly, a control mouse receiving c-Jun_CaffCAR T cells but no Raji tumor cells (to be able to assess potential background BLI signal caused by the pumps) also developed a large liver nodule (Fig. 6A). Even though this was only one mouse and therefore interpretation requires caution, this further supports the conclusion that this effect is CAR-independent.

In line with the emergence of white nodules, we found markedly elevated T cell numbers in the livers of c-Jun_CaffCAR T cell-treated mice, independent of caffeine (Fig. 6B and Suppl. Fig. 5E), contrasting with CaffCAR T cells whose expansion was caffeine-dependent. Similar hyperproliferation was observed in blood, spleen and bone marrow (Fig. 6B and Suppl. Fig. 5E), suggesting that c-Jun overexpression induces systemic T cell expansion in a CAR-independent manner.

## Discussion

To address the poor controllability of CAR T cells in patients, we engineered CaffSwitches that offer multiple crucial advantages including (i) defined heterodimerization that is highly dependent on the presence of caffeine, (ii) the use of small protein domains (119 amino acids), (iii) an appropriate affinity of ∼20 nM between the two dimerizing nanobodies, (iv) efficient function in the reducing cytoplasmic environment, (v) rapid ON- and OFF-kinetics and (vi) virtually absent leakiness. Finally, we demonstrated that these switches can be used to generate CaffCARs, which are highly responsive to caffeine levels reached by drinking one cup of coffee. Since CaffSwitches are modular, we were able to generate CaffCARs with different antigen specificities and costimulatory domains.

When using caffeine-responsive switches *in vivo*, unintended background activation may be caused by leakiness in the absence of caffeine, as well as responsiveness to theobromine (contained in chocolate) or traces of caffeine. Since all these effects were observed with the original homodimeric switch, we addressed these issues during the engineering process: First, by using rather high (10 µM) caffeine concentrations in positive selections, we were able to reduce activation of the CaffSwitches by trace levels of caffeine. Second, to avoid triggering upon chocolate consumption, we excluded the theobromine-responsive variant acVHH-M2. Third, we performed rigorous negative selections in the absence of caffeine, which resulted in tightly controlled switches with virtually absent leakiness. This is also reflected *in vivo* by the lack of tumor control of c-Jun_CaffCAR T cells in the absence of caffeine, despite their massive expansion (Fig. 5D and 6B).

Small molecule-regulated switches have the potential to address multiple limitations of CAR T cells^54^. For example, it has been demonstrated that transient rest prevents T cell exhaustion and thereby improves T cell potency and persistence^13,14^. In line with these previous studies, we showed that ON/OFF cycles with CaffCAR T cells reduces exhaustion and promotes CAR T cell expansion. Moreover, in the event of severe toxicities such as cytokine release syndrome (CRS), neurotoxicity or on-target/off-tumor toxicity (OTOT)^55,56^, the ability to control the activity of this living drug would be a key advantage. One example for OTOT is long-term B cell aplasia linked to the desired persistence of CD19-reactive CAR T cells^8–10^. Drug-regulated CAR T cells could be inactivated after successful therapy of B-cell lymphomas, where long-term persistence may be dispensable^57^, to enable return to normal B cell responses. In the case of solid tumors, OTOT against more critical tissues can even be fatal^58^. These toxicities are expected to be even more severe and prolonged with next generation CAR T cells engineered for improved potency and persistence. Indeed, a recent phase I study with c-Jun overexpressing CAR T cells needed to be halted due to dose-limiting toxicities, including grade 4 CRS that did not respond to tocilizumab or corticosteroids^59^. Importantly, we demonstrate that the activity of such boosted, c-Jun overexpressing CAR T cells can be tightly controlled by integrating the CaffSwitch into the CAR.

One unexpected finding in this study was the formation of white nodules in the livers of most mice treated with c-Jun overexpressing CaffCAR T cells. In addition, we observed massively increased T cell numbers in the liver, spleen, bone marrow and blood of these mice, hinting towards c-Jun-induced CAR T cell hyperproliferation. Strikingly, both nodule formation and T cell expansion seemed to be independent of caffeine-induced CAR signaling. Although the formation of nodules has not been reported before, Lynn et al. observed strong expansion of c-Jun overexpressing CAR T cells in tumor-free mice^11^, which is in line with our findings that this effect is largely CAR-independent. While this interesting phenomenon clearly warrants further investigation, this is outside the scope of the present study.

When compared with small molecule drugs used in other molecular switches, caffeine offers a range of highly advantageous properties. Caffeine is a well-characterized drug extensively studied in humans. It is readily available, inexpensive and known to be non-toxic at reasonable concentrations even when consumed over decades^32,33^. While for pregnant women lower caffeine consumption is recommended, doses up to 2-3 cups of coffee per day are still considered safe by the Food and Drug Administration (FDA) and the European Food Safety Authority (EFSA)^60,61^. Due to its well-established safety profile, caffeine is even routinely used to treat apnea of prematurity in infants^34,35^. Moreover, due to its efficient distribution in the human body, including its ability to cross the blood-brain-barrier, caffeine enables the regulation of CaffCARs in various tissues and organs^31^. Its half-life of ∼5 h in humans^31^ is ideally suited for efficient deactivation of CaffCARs *in vivo* within a reasonable time frame.

Caffeine plasma levels in individuals avoiding caffeine consumption were shown to be below 20 nM (limit of quantification)^62^. Importantly, at these low concentrations, our CaffCAR T cells were completely inactive. On the other hand, drinking one cup of coffee results in plasma levels of ∼10 µM caffeine^51,52^, which is sufficient to trigger full activation of CaffCAR T cells. Moreover, kinetic experiments showed that the CaffCARs are rapidly turned on and off after administration and removal of caffeine, respectively, demonstrating that the activation state of the CaffCARs rapidly adapts to the available caffeine concentration.

Summing up, we engineered novel CaffSwitches which heterodimerize in a caffeine-dependent manner and show a number of highly beneficial properties. We demonstrate that CaffSwitches can be used to generate CaffCAR T cells which efficiently respond to caffeine concentrations found in human plasma after drinking one cup of coffee. In addition to split CARs generated in the present study, caffeine-responsive switches also enable the regulation of transcription factors and cytokine receptors in T cells, as demonstrated in a companion paper by Scheller et al. Therefore, we anticipate that CaffSwitches will be highly valuable tools for the regulation of various synthetic components in CAR T cells and other cellular therapies.

## Acknowledgements

This work was supported by the Federal Ministry for Digital and Economic Affairs of Austria, and the National Foundation for Research, Technology and Development of Austria to the Christian Doppler Research Association (Christian Doppler Laboratory for Next Generation CAR T Cells) and by private donations to the St. Anna Children’s Cancer Research Institute (Vienna, Austria). This research was also funded in part by the Austrian Science Fund (FWF) [10.55776/P34832; W1224–Doctoral Program on Biomolecular Technology of Proteins–BioToP; EFP 45, Devising Advanced TCR-T cells to eradicate OsteoSarcoma, DART2OS)] and the Rete ACC - Sviluppo successivo per il progetto CAR-T rete Oncologica: ponte alla traslazione clinica (RCR-2023-23684268). E.S. and M.C.B. are recipients of DOC Fellowships of the Austrian Academy of Sciences at the St. Anna Childreńs Cancer Research Institute (#26323 and #25905). Research at the IMP is supported by Boehringer Ingelheim and the Austrian Research Promotion Agency (headquarter grant FFG-852936). MS is a member of the Boehringer Ingelheim Discovery Research global post-doc program. E.M. is supported by the Boehringer Ingelheim Fonds (BIF). Research in the van der Veeken laboratory is supported by Boehringer Ingelheim, the Austrian Science Fund (FWF, 10.55776/PAT4163824) and the European Research Council (ERC) under the Horizon 2020 research and innovation program (ERC-2023-STG 101116251). We further acknowledge the CCRI FACS Core Unit for providing cell sorting and analysis services, the Connective Base GmbH, BOKU Core Facility Biomolecular & Cellular Analysis and BOKU Core Facility Mass Spectrometry for providing flow cytometers, and MS equipment, and supporting the project. All schematic figures were created with Biorender.com.

## Data availability

Source data will be provided with this paper and deposited in a publicly available repository before final publication.

## Author contributions

E.S., B.S., M.L. and M.W.T. conceived the study. E.S., B.S., D.E., H.B., G.D., M.S., K.M., M.C.B. and C.U.Z. performed experiments and acquired data. E.S., B.S., D.E., H.B., G.D., M.S., E.M., A.M.-L., F.E., B.E., J.M., J.V., A.R., J.Z., E.M.P., C.U.Z., M.L., M.W.T. designed experiments. E.S., D.E., H.B., G.D., M.S., K.M., C.U.Z., M.L., M.W.T., analyzed data and interpreted results. J.S. expressed and purified the soluble stabilized CD19 protein. D.M. and A.U. developed the method for mass spectrometric quantification of caffeine in mouse plasma, performed the measurements and data analysis. M.L. and M.W.T. supervised the work. E.S. and M.W.T. wrote the manuscript. All authors edited and approved the manuscript.

## Materials and Methods

### Recombinant expression of proteins

The different acVHH variants were expressed recombinantly as soluble proteins fused to an N-terminal hexahistidine tag using the pET21+ vector. Briefly, *Escherichia coli* cells (Rosetta cells) were transformed with the respective sequence verified plasmids using heat shock transformations. After overnight incubation in lysogeny broth (LB) medium supplemented with ampicillin (100 µg/mL) at 37 °C, 180 rpm, cultures were set to an optical density at 600 nm (OD_600_) of 0.2 and further incubated at 37 °C. Induction of transgene expression was initiated by addition of 1 mM of isopropyl β-D-1-thiogalactopyranoside (IPTG) when the OD_600_ reached 1, followed by overnight incubation at 20 °C. Cells were harvested the next day by centrifugation (5000 g, 20 min, 4 °C). The pellet was resuspended in sonication buffer (50 mM phosphate buffer pH 7.5 containing 300 mM NaCl, 3 % glycerol, 1 % Triton X-100) and the cells were lysed by sonication (2 min, pulse: 1,0 – 1,0 seconds, 90 % amplitude) on ice. The cell debris were removed by an additional centrifugation step (20 000 g, 30 min at 4 °C). The supernatant containing the soluble proteins was supplemented with 10 mM imidazole and filtered through a 0.45 µM filter (Duropore). Hexahistidine-tagged proteins were purified by immobilized metal affinity chromatography using a HisTrap FF column (Cytiva) connected to a Bio-Rad purifier system (Bio-Rad NG-C chromatography system). After loading of the supernatant and washing steps (50 mM phosphate buffer pH 7.5 with 500 mM NaCl), elution of the pure proteins was carried out by applying a linear imidazole gradient (from 5 % to 100 % of a buffer containing 50 mM phosphate, pH 7.5, 500 mM NaCl and 500 mM imidazole). Absorbance at 280 nm was detected and the fractions containing a protein of the right molecular weight (as analyzed via SDS-PAGE) were pooled. Buffer exchange to PBS (Thermo Scientific) was performed overnight at 4 °C via dialysis using a 10 kDa SnakeSkin dialysis tubing (Thermo Scientific). Purified proteins were concentrated via centrifugation steps in Amicon Ultra-15 10 K centrifugal filters (Merck Millipore). A fraction of acVHH-V106D was labelled with biotin using the EZ-Link Sulfo-NHS-LC-LC-Biotin kit (Thermo Scientific) for use in magnetic bead selections.

### Size Exclusion Chromatography (SEC)

Protein aggregates or unbound biotin molecules were removed via preparative size exclusion chromatography. After a centrifugation step (16 000 rpm for 3 min) the supernatant was loaded onto a HiLoad Superdex 75 pg column connected to a Bio-Rad NG-C chromatography system. PBS was used as running buffer and the fractions containing monomeric proteins based on absorbance at 280 nm were pooled, concentrated as mentioned above and stored at −80 °C.

### Yeast surface display library preparation and screening for binders

For the first library, an acVHH gene fragment with 8 NNK randomized positions (amino acids 52-59) was ordered (Twist) and additionally submitted to error-prone PCR (ep-PCR) mutagenesis using the GeneMorph II – random mutagenesis kit (Agilent) with the following parameters: 100 ng template DNA, 20 cycles.

For the second library, inserts coding for acVHH-M1 ΔAA, -M1 ΔAV, -M1 ΔVA, and -M1 ΔWP were amplified by PCR, pooled in an equimolar ratio, and subjected to error-prone PCR as described above.

Subsequent insert and vector (pCTCON2V) preparation steps as well as electroporation of the library in electrocompetent *S. cerevisiae* cells (strain EBY100, ATCC) were prepared according to the protocol published by Chen et al.^63^ Library diversities were calculated by plating serial dilutions of the cells on selective SD-CAA plates after electroporation and bead selections, and allowing for growth of transformants.

After electroporation, the yeast library was cultivated to an OD_600_ of 1 in SD-CAA (20 g/L glucose, 6.7 g/L yeast nitrogen base, 5 g/L casamino acids, 11.85 g/L sodium citrate dihydrate and 7.4 g/L citric acid monohydrate) at 30 °C, 180rpm. Protein expression on the yeast surface was induced by switching the medium to SG-CAA (2 g/L glucose, 20 g/L galactose, 6.7 g/L yeast nitrogen base, 5 g/L casamino acids, 10.2 g/L disodium hydrogen phosphate and 4.82 g/L sodium phosphate monobasic) and subsequent incubation overnight at 20 °C.

Bead selection rounds were performed as described previously^63^ with magnetic streptavidin-coated Dynabeads (Life Technologies) loaded with biotinylated acVHH-V106D. Briefly, two cycles of bead selections were performed per engineering campaign with alternating positive and negative selections. In positive selections, cells displaying acVHH variants that could bind to acVHH-V106D-coupled beads in the presence of 10 µM caffeine were selected, whereas binding to bare beads was excluded via negative selections. The reduced library was further enriched by fluorescence activated cell sorting (FACS). The yeast library was incubated with 500 nM soluble acVHH-V106D for 1 h at 4 °C while shaking in the presence (positive selections) or absence (negative selections) of 10 µM caffeine (Sigma) in PBSA (PBS supplemented with 15 µM bovine serum albumin). For all *in vitro* experiments, a fresh stock of 10 mM caffeine was prepared (dissolved in water and sonicated for 10 min) which was further diluted in PBSA to the desired concentration. Biotinylated and non-biotinylated hexahistidin-tagged acVHH-V106D were used in different selection rounds to prevent selection of biotin specific binders. After a washing step the cells were stained for 20 min at 4 °C with shaking with either Penta-His-Alexa Fluor 647 (Qiagen) or Streptavidin-Alexa Fluor 647 (Invitrogen) to assess binding to acVHH-V106D, as well as anti-HA-Alexa Fluor 488 (clone 16B12, Biolegend) or anti-c-myc-Alexa Fluor 488 (9E10, Thermofisher) to assess display (anti-HA) or full-length display (anti-c-myc) of the acVHH variants. After a final washing step with PBSA, cells were sorted using a SONY Sorter SH800.

After the last selection round, plasmids were obtained from the yeast libraries using a Zymoprep yeast plasmid miniprep kit II (Zymo Research). The resulting DNA pool was electroporated into *E. coli* 10-beta electrocompetent cells (NEB) according to the supplier’s protocol, followed by sequencing of individual clones (Microsynth). Plasmids of individual acVHH-variants of interest were subsequently transformed into the *S. cerevisiae* strain EBY100 with the Frozen-EZ Yeast Transformation II Kit (Zymo Research).

For *K*_D_ and *EC*_50_ values based on yeast surface display, the geometric mean fluorescence intensity (gMFI) binding values were normalized and fitted to a 1:1 binding interaction model^64^. Negative values were set to 0.

### Mammalian cell culture

Buffy coats from healthy donors were purchased from the Austrian Red Cross, Vienna, Austria. Primary human T cells were purified by negative selection using the RosetteSep Human T cell Enrichment kit (STEMCELL Technologies) and cryopreserved in RPMI-1640 GlutaMAX medium (Thermo Scientific) supplemented with 20 % FCS and 10 % DMSO (Sigma-Aldrich). T cells were cultivated between 0.3 - 2×10^6^ cells/mL in AIMV medium (Thermo Scientific) supplemented with 2 % Octaplas (Blutspendezentrale Wien), 2.5 % Hepes (Pan Biotech), 1 % glutamine (Gibco) and 200 U/mL recombinant human IL-2 (Peprotech). Raji, Jurkat, LNCaP (gift from Dr. Sabine Strehl, Dr. Michael Dworzak, and Dr. Thomas Lion, respectively, CCRI, Vienna) and PC3 cells (ATCC) were transduced with a lentiviral vector encoding ffLuc and enhanced GFP and maintained in RPMI-1640 GlutaMAX supplemented with 10 % FCS, 1 % penicillin–streptomycin (Thermo Scientific). Additionally, PC3-GFP/ffLuc cells stably expressing PSMA were generated in house. Lenti-X 293T cells (Takara) were maintained in DMEM (Thermo Scientific) supplemented with 10 % FCS.

### *In vitro* transcription and mRNA electroporation

After PCR amplification of the transgene, 200 ng of purified product were transcribed *in vitro* using the mMessage mMachine T7 Ultra Kit (Ambion) according to the manufactureŕs instructions. The resulting mRNA was purified with the RNeasy Kit (Qiagen). Cells were electroporated with 5 µg mRNA per chain using 4 mm cuvettes (VWR) and the Gene Pulser (Bio-Rad) with the following parameters: square wave protocol, 500 V, 3 ms for Jurkat T cells or 5 ms for primary human T cells. Electroporated cells were recovered in warm cell growth medium and incubated at 37 °C.

### Intracellular protein complementation assay

Intracellular binding of proteins was determined with the Nano-Glo® Luciferase Assay System (Promega). The acVHH variants were N-terminally fused to an extracellular globular domain to assess expression (truncated EGFR or truncated HER2), and C-terminally fused to a subunit of the split Nanoluc (NanoBiT® system, Promega). To assess the heterodimerization of proteins, acVHH-V106D was always fused to truncated EGFR and 11S NanoBiT® “Large BiT”, while acVHH-M1, -M2, dfM1-1 and dfM1-2 were fused to truncated HER2 and 114 NanoBiT® “Small BiT”. 16 h after electroporation, 100 000 cells expressing the proper chains were cultured in the presence of the appropriate compound for 30 min at 37 °C. The Nano-Glo reagents were reconstituted as suggested by the manufacturer and added to the cells right before measuring luminescence (ENSPIRE plate reader). For titrations in Jurkat T cells, *EC*_50_ values were calculated by fitting the RLU binding values to a 1:1 binding interaction model^64^ after subtracting the background signal (Jurkat T cells not electroporated but cultured with the highest concentration of the appropriate compound).

### Lentivirus production

Lenti-X 293 T cells (Takara) were co-transfected by second generation viral packaging plasmids pMD2.G and psPAX2 (Addgene plasmids #12259 and #12260, respectively; gifts from Didier Trono) and either a third generation puromycin selectable pCDH expression vector (System Biosciences) for chain II (CD3z – signaling) genes, or a pCDH vector with the puromycin gene removed and a stop codon introduced after the chain I (αCD19 or αPSMA) transgene. For the cJUN_CaffCAR, c-Jun was added on the same vector as chain II as follows: c-Jun_2A_chain II. Transfections were carried out using the PureFection Transfection Reagent (System Biosciences) according to the instructions provided by the manufacturer. Viral supernatants were collected on day 2 and 3 after transfection and concentrated 50- or 100-fold (for chain II or chain I, respectively) using the Lenti-X Concentrator (Takara) according to the manufacturer’s instructions. Viruses were resuspended in AIMV medium supplemented with 2 % Octaplas, 2.5 % Hepes, 1 % glutamine and 200 U/mL IL-2 and stored at −80 °C.

### CAR-T cell manufacturing via lentiviral transduction

On day 0 primary human T cells were thawed and activated with Dynabeads Human T-Activator αCD3/αCD28 beads (Thermo Scientific) at a 1:1 bead-to-cell ratio. On day 1, 100 µL T cells (1 x 10^6^ cells/mL) were seeded in RetroNectin (Takara) coated 96 well plates and co-transduced with the vectors encoding chain I and II by adding thawed viral supernatant at a 2:1:1 final dilution (cells:chain I:chain II). T cells expressing chain II were selected by adding 1 µg/mL puromycin (Sigma-Aldrich) on day 3. CAR T cells were used 8 to 12 days after transduction for cytokine secretion experiments, and between 15 to 25 for cytotoxicity assays. For the exhaustion experiments (Figure 4) and mouse experiments (Figure 5 and 6), cells were sorted for chain I expression 4 days after puromycin expression (chain II selection) by using a soluble stabilized CD19 antigen conjugated with AF647 made in house^65^.

The αCD19-containing chains (chain I of the split CARs or CD19 control CAR) and the αPSMA-containing chains (chain I of the split CARs or PSMA conventional CAR) include a FLAG tag, the CD3z signaling chains (chain II) either have an N-terminal StrepII tag or are fused to mCherry, which were used to detect CAR expression and determine CAR^+^ fractions to set up proper E:T ratios.

### Flow cytometric analysis of CAR expression and cell counting

Nonspecific antibody binding was prevented by preincubating the cells for 10 min at 4 °C in FACS buffer (PBS with 0.2 % human albumin (CSL Behring) and 0.02 % sodium azide (Merck KGaA)) supplemented with 10 % (v/v) human serum. Afterwards cells were incubated with the antibodies for 25 min at 4 °C in FACS buffer, washed twice, acquired on a BD FACSymphony A3 Cell Analyzer or BD LSR Fortessa (BD Biosciences) and analyzed with the Flowjo Software. Non-transfected or non-electroporated cells served as negative controls. The following antibodies were used: αFLAG (APC or PE; clone L5; BioLegend), αStrepII (FITC; clone 5A9F9; Genescript), αCD19 (APC or PE; clone HIB19, BioLegend), αCD3 (BV605; clone OKT3, Biolegend), αCD8 (BUV496; clone RPA-T8, BD Horizon), αCD4 (PerCP, clone OKT4; Biolegend), αCD39 (PE-Vio770; clone REA739, Miltenyi Biotec), αPD1 (APC; clone REA1165, Miltenyi Biotec), αLAG3 (APC_Cy7; clone 11C3C65, Biolegends), αCD62L (PEVio770; clone REA615, Miltenyi Biotec), αCD45RA (APC; clone REA562, Miltenyi Biotec). The following isotype controls were used: Mouse IgG1 κ (BV421 and APC_Cy7; clone MOPC-21, Biolegends) and REA TM (PEVio770 and APC; clone REA293, Miltenyi Biotec). The following fixable viability dyes were used: BD Horizon™ Fixable Viability Stain 440UV and 780 (BD Biosciences).

When FACS panels included multiple brilliant violet-conjugated antibodies, mixes and incubations were carried out in brilliant stain buffer (BD Horizon). When FACS panels included a fixable viability dye, serum-free washing buffer (plain PBS) was used.

Accurate cell counts were obtained by adding 10 µL of AccuCheck Counting Beads (Thermo Scientific) to the cells after staining them and normalizing cell counts to the number of beads acquired.

### Cytotoxicity assays

All cytotoxicity assays were luciferase-based and carried out in RPMI without phenol red (Thermo Scientific) supplemented with 2 % Octaplas and 1 % Pen/Strep. Each experiment was set up with 3 technical replicates per condition. Target cells expressed GFP and luciferase.

For the Jurkat T cells killing assay (Figure 2A), expression of the CARs in primary human T cells as well as the CD19 antigen (SinoBiological, RefSeq BC006338) in Jurkat T cells was achieved via mRNA electroporation (5 mg). Mock primary human T cells (not electroporated) served as control. 6 h after electroporation 30 000 effector T cells were co-cultured with target cells in 96 wells U bottom plates at a 2:1 E:T ratio in phenol-free media containing medium only for controls or the appropriate compound: 0.5 µM AP21967 for FKBP/FRB dimerization (Takara) or 50 µM caffeine. After a 4 h co-culture, 150 µg/mL luciferin was added, and luminescence was measured 20 min afterward with an ENSPIRE Multimode plate reader (Perkin Elmer). Specific lysis was calculated according to the formula:

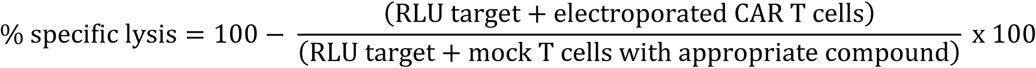

For the CaffCAR experiments (Figure 2B to 2E and Figure 3), CAR expression was achieved with lentiviral transductions. CAR^+^ T cells were co-cultured with 10 000 Raji target cells at an E:T ratio of 2:1 overnight at 37 °C in the presence of the appropriate compound. Mock T cells were added to reach the same total number of T cells among the different conditions and account for different CAR positive fractions. The next day luminescence was measured as described above and specific lysis was calculated according to the formula:

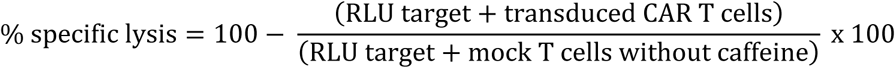

### Long-term co-culture experiment

CaffCAR T cells were transduced as indicated above, including selecting for chain II expressing cells via puromycin selection on day 3. Beads were removed on day 6 and cells expressing chain I were sorted on day 7 with a BD FACSAria Fusion by staining with a soluble stabilized CD19 antigen conjugated with AF647 made in house^65^. Mock T cells used as control had also their beads removed on day 6 and were sorted for total cells on day 7.

Co-cultures with Raji-GFP/ffLuc target cells were set up in triplicates for each donor in 2 mL (E:T 1:1, 0.5 Mio each) in 12 well plates in the presence (CaffCAR ON) or absence of 30 µM caffeine (CaffCAR OFF and mock) (day 0). Cells were counted every 2-3 days via flow cytometry using AccuCheck Counting Beads (Thermo Scientific) and split accordingly. The percentage of Raji target cells in each well was assessed based on GFP expression. On day 4 the triplicates CaffCAR ON co-cultures were washed, split into two each, and caffeine was either added (CaffCAR ON) or not (CaffCAR pause). To alternate between ON- and OFF-states in the CaffCAR pause co-cultures, the entire co-cultures were either centrifuged and washed to remove all trace of caffeine (ON to OFF), or caffeine was simply added (OFF to ON). On day 7, new Raji tumor cells were added to the CaffCAR ON and CaffCAR pause cultures to reach again an E:T of 1:1. Flow cytometric stainings for exhaustion markers were performed on day 17, 28 and 37.

### Cytokine release assays

Co-culture supernatants of triplicate wells from the experiments described above were pooled and frozen for the analysis of secreted cytokines. IFN-γ and IL-2 secretion was quantified by enzyme-linked immunosorbent assay (ELISA) using the ELISA MAX Deluxe Set Human IFN-γ and ELISA MAX Deluxe Set Human IL-2 respectively (Biolegend) according to the manufacturer’s instructions. The data was acquired on the ENSPIRE Multimode plate reader (Perkin Elmer) and the analysis was carried out using the GainData® online tool (Arigo), excluding all values reaching the standard curve’s plateau.

For OFF kinetics, CaffCAR T cells were cultured in the presence of 30 µM caffeine 48 h prior to the assay. At the indicated time points (30-24-8-4-1 h prior to start of the co-culture assay), cells were washed to remove the drug and cultured in plain medium. A 24 h co-culture with Raji (E:T of 2:1) was set up, followed by analysis of IFN-γ and IL-2 in the supernatant. CaffCAR cells that were always in the presence of caffeine were washed once and caffeine was added (ON) or not (0 h) in the co-culture. Mock T cells as well as CaffCAR cells that were never cultured with caffeine (OFF) were used as negative controls.

For ON kinetics, 30 µM caffeine was added in the CaffCAR T cell growth medium at the indicated time point (24-20-4-2 h) before the assay. CAR^+^ T cells were co-cultured at an E:T ratio of 2:1 with Raji cells for 4 h and IFN-γ concentration in the supernatant was measured. For the 0 h time point caffeine was only added in the co-culture medium.

### *In vivo* experiments

NOD.Cg-Prkdc^scid^ Il2rg^tm1WJI^/SzJ (NSG, The Jackson Laboratory) mice were kept at the Core Facility Laboratory Animal Breeding and Husbandry of the Medical University of Vienna (CFL) under specific pathogen-free conditions according to FELASA recommendations (Himberg, Austria). Animal experiments have been carried out either at the CFL (Vienna) or the Institute for Molecular Pathology (IMP, Vienna). All procedures were approved by the Magistratsabteilung 58, Vienna, Austria (GZ: MA 58 – 1390324-2024-19) and the Austrian Federal Ministry of Education, Science and Research (GZ 66.009/0243-V/3b/2019) and were performed according to the guidelines of FELASA and ARRIVE. Primary human T cells were lentivirally transduced for CAR expression and expanded for 12 - 13 days. On day 3, cells expressing the CD19 CAR or the chain II of the CaffCAR were selected by puromycin selection (1 µg/mL). At day 7, cells expressing the CD19 CAR or the chain I of the CaffCAR were sorted with a BD FACSAria Fusion by staining with a soluble stabilized CD19 antigen conjugated with AF647 made in house^65^. Mock T cells used as control were sorted for total single cells. 0.5 x 10^6^ Raji cells expressing GFP and ffLuc were injected i.v. into the tail veins of NSG female mice (10 – 22 weeks old). 3 x 10^6^ or 5 x 10^6^ CAR positive T cells were injected i.v. three or five days later (as indicated). When caffeine was administered via the drinking water, the small molecule was dissolved to 7 g/L in water, sonicated 15 minutes (Sigma Aldrich), sterile-filtered through a 0.2 µm membrane filter (Nalgene), and diluted to 0.7 g/L in the sterile mouse water bottles. Drinking bottles were exchanged three times per week. When caffeine was administered via osmotic pumps, caffeine was dissolved to 250 g/L in 2.5 M sodium benzoate (Sigma Aldrich), used to increase the aqueous solubility of caffeine^66^. The caffeine solution was then sterile-filtered using a 0.2 µm membrane filter and filled into ALZET Mini-Osmotic Pumps (model 2004, 200 µL, release of 0.25 µL/hour for up to 28 days) according to the supplier’s instructions. To avoid any bioluminescence background, ALZET Flow Moderator Blue (ALZET, 0002489) was used together with the osmotic pumps. For mice not receiving caffeine, pumps containing only 2.5 M sodium benzoate were also prepared as control. All pumps were filled one day before implantation and incubated in 0.9 % sodium chloride solution at 37 °C overnight.

One day after tumor cell injection, the mice were shaved and received intraperitoneal (i.p.) injections of 5 mg/kg carprofen (Rimadyl, Zoetis) pre-emptively and 12 h after surgery. They were then anesthetized with isoflurane (Vetflurane 1000 mg/g, Virbac), pumps were implanted subcutaneously in the back of the mice and their wounds were clipped. In the experiment shown in Figure 5C, 5D and Figure 6, all mice were sacrificed at the same time point (day 22 after tumor injection).

### *In vivo* bioluminescence imaging

Bioluminescence imaging (BLI) was performed at the IMP using the IVIS Spectrum imaging system (PerkinElmer). Mice were anesthetized with isoflurane (Vetflurane 1000 mg/g, Virbac) and received

i.p. injections of D-luciferin (150 mg/kg body weight, Goldbio). After 5 min, mice were transferred to the IVIS Imaging System and bioluminescence was measured in medium binning mode with an automatic acquisition time. Tumor growth was monitored every 3–4 days. Living Image^TM^ software (v.4.8, Revvity) was used to analyze the data and extract total photon fluxes. Since groups were randomized based on tumor burden on d0 and mice in the same cage got different T cell treatments, the pictures were cropped to reconstitute Figure 5C.

### Mouse organ processing

Blood was collected via tail veins into EDTA tubes. Bone marrow single-cell suspensions were obtained by flushing two femurs with 15 mL PBS and subsequent filtering through a 70 µm cell strainer. Spleen samples were disrupted and passed through 70 µm filters twice. Liver cells were first dissociated using a liver dissociation kit (Miltenyi Biotec) due to the presence of solid nodules in some of them. They were then disrupted and filtered once through 70 µm filters. The liver single-cell suspensions were subjected to a 33.75 % Percoll gradient centrifugation to separate lymphocytes from hepatocytes. All single-cell suspensions were subjected to multiple washing steps in PBS and lysis of red blood cells with ACK Lysing Buffer. Finally, cells were blocked with 5 % Octaplas and a fraction of the volume was stained in 50 µL antibody mix (anti-CD3-BV605, final dilution 1:50; anti-Flag-PE 1:100; anti-CD19-APC, 1:100; anti-StrepII-FITC, 1:25; fixable viability dye 780, 1:1000), washed twice and measured on a BD LSR Fortessa together with 10 µL AccuCheck Counting Beads (Thermo Scientific).

### Mass spectrometric analysis of caffeine concentrations in mice plasma

Whole blood was collected from mice in EDTA tubes and plasma was harvested by centrifuging at 2 000 g for 15 min and collecting the supernatant.

Caffeine powder (Sigma Aldrich; PN: C0750) was used for the preparation of calibration standards. ^13^C_3_-labeled caffeine (Sigma Aldrich; PN: C082) served as the internal standard and was added in equal volumes to all calibration standards and samples. Calibration standards were spiked with negative plasma from control mice to achieve matrix matching. LC-MS grade acetonitrile (Thermo Scientific; PN: A956-1) was added to both standards and samples to precipitate plasma proteins. The mixture was cooled at 4 °C for 15 minutes, followed by centrifugation at 21 300 g for 15 minutes. The resulting supernatant was diluted with an equal volume of LC-MS grade water prior to injection.

Chromatographic separation was performed on a ZORBAX SB-Aq C18 column (Agilent, 2.1*50 mm, 1.8 µm) using 0.1 % formic acid in LC-MS grade water as the aqueous mobile phase (A) and 0.1 % formic acid in LC-MS grade acetonitrile as the organic mobile phase (B). Initial conditions were set to 5 % B (0.5 min), ramped linearly to 80 % B over 2.5 min, held at 80 % B for 0.9 min, and then returned to baseline conditions for 2 min. The total run time was 6 minutes with a flow rate of 300 µL/min and a column oven temperature of 40 °C. The injection volume was 5 µL.

For the quantification of caffeine in plasma from mice that received caffeine via drinking water (n = 9), detection was carried out using a TSQ Vantage triple quadrupole mass spectrometer (Thermo Scientific) equipped with a H-ESI source in positive ion mode. The ion source parameters were as follows: vaporizer temperature, 275 °C; ion transfer tube temperature, 275 °C; auxiliary gas pressure, 15 arbitrary units; sheath gas pressure, 40 arbitrary units; ion sweep gas pressure, 0 arbitrary units; spray voltage, 3300 V. Selected reaction monitoring (SRM) was utilized to quantify caffeine, with the following transitions: 195.1 *m/z* → 138.0 *m/z* for caffeine and 198.1 *m/z* → 140.0 *m/z* for ^13^C_3_-labeled caffeine. The collision energy for the quantifier transitions was 18 eV, and the collision gas pressure was 1.5 mTorr. Data analysis was performed using TraceFinder 5.2.

For the quantification of caffeine in plasma from mice that received caffeine via osmotic pumps (n = 10), detection was performed using an Orbitrap IQ-X Tribrid LC-MS^n^ mass spectrometer (Thermo Scientific) equipped with a H-ESI source in positive ion mode. The ion source parameters were set as follows: vaporizer temperature, 325 °C; ion transfer tube temperature, 275 °C; auxiliary gas pressure, 10 arbitrary units; sheath gas pressure, 50 arbitrary units; ion sweep gas pressure, 1 arbitrary unit; spray voltage, 3200 V. The Orbitrap resolution was set to 120 000, with an RF Lens setting of 60% and a maximum injection time of 246 ms. RunStart EASY-IC was utilized for internal mass calibration. A targeted selected ion monitoring method (tSIM) method was employed for both caffeine (195.0877 *m/z*) and ^13^C_3_-labeled caffeine (198.0977 *m/z*), with both ions being multiplexed. Data analysis was performed using Skyline 21.2.0.425.

### Statistical analysis

Statistics were performed with GraphPad prism software version 9 for Windows (GraphPad Software Inc) and are indicated under each figure. *EC*_50_ values based on lysis or cytokine release were calculated with the [Agonist] vs response function of GraphPad. Statistics for cytokines secretion, as well as T cell and CAR T cell numbers in mouse organs were performed on log-transformed values.

