## Supplementary material for "Caffeine-regulated molecular switches for functional control of CAR T cells *in vivo*": Suppl. Information

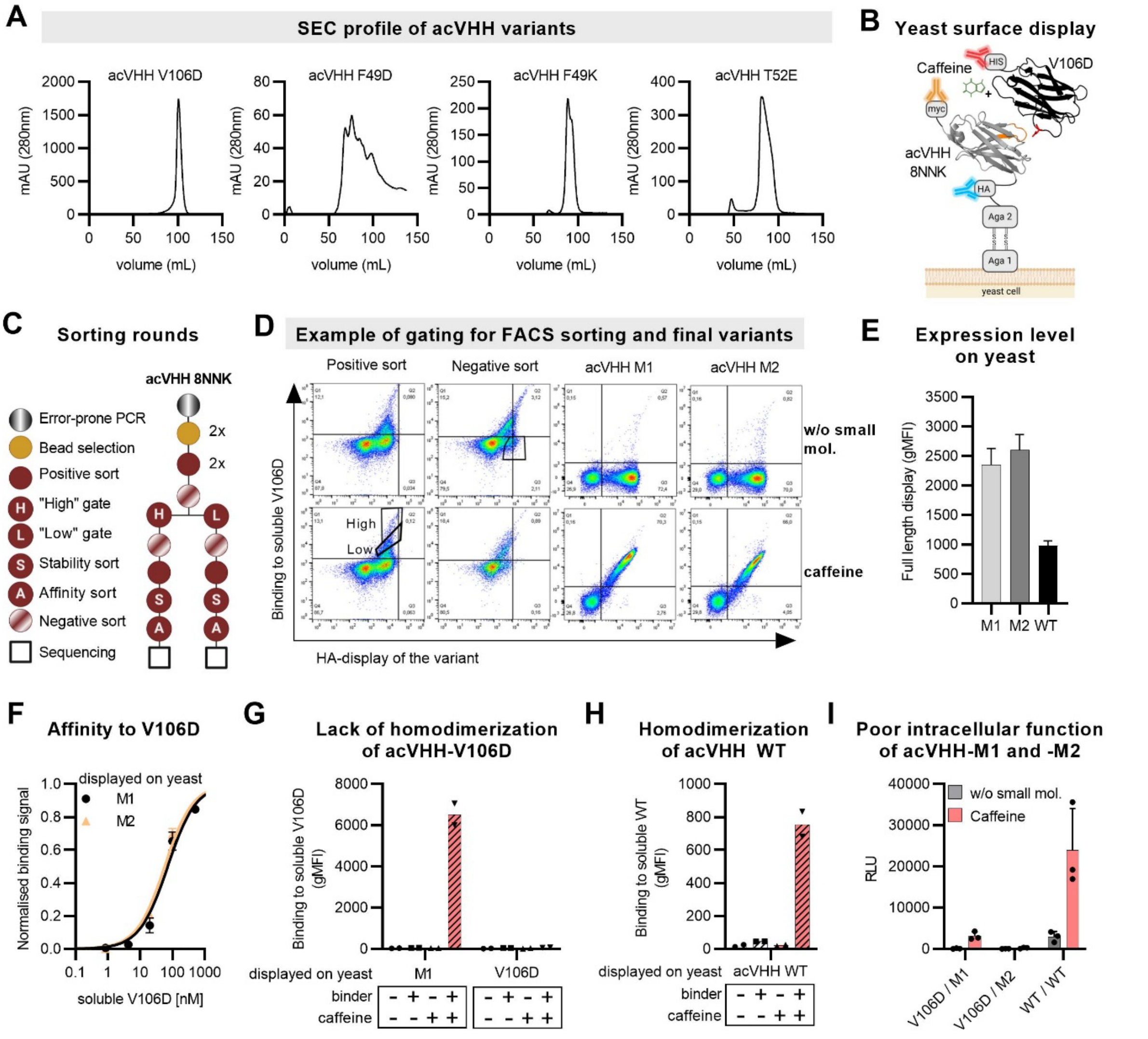


**Supplemental Figure 1: Engineering of caffeine-regulated heterodimeric switches.**

**(A)** Size exclusion chromatography (SEC) profiles of acVHH variants containing single point mutations as indicated.

**(B)** Schematic of the yeast surface display platform used for the engineering process. The nanobodies of interest (for example the acVHH-8NNK library) are expressed extracellularly and anchored to the yeast surface via its fusion partner Aga2. HA- and c-myc tags allow for analysis of surface expression levels. His-tagged acVHH-V106D is added solubly and detected with an anti-His antibody.

**(C)** Detailed schematic of the selection strategy. In positive sorts, variants were selected for binding to soluble acVHH-V106D in the presence of caffeine, whereas in negative sorts, variants were enriched for non-binding to acVHH-V106D in the absence of caffeine. The library was split into “high” and “low” gates to enrich more stringently for variants with a high binding signal while also keeping the ones with a “low” binding signal.

**(D)** Expression on the yeast surface (measured via HA-tag) is shown on the x-axis and binding to acVHH-V106D is shown on the y-axis. In the left column, flow cytometry sorting gates of a representative positive sort are shown, where the cells that bound to acVHH-V106D in the presence of caffeine were sorted in “high” or “low” gates. In the second column, a representative negative sort is shown, where cells displaying acVHH variants that did not bind to acVHH-V106D in the absence of caffeine were sorted.

**(E)** Full-length display levels of acVHH variants on yeast cells, as measured via the c-myc-tag fused to the C-terminus of the nanobodies (mean ± SD, n=3).

**(F)** Binding of the yeast-displayed acVHH mutants M1 and M2 to increasing concentrations of soluble acVHH-V106D was assessed in the presence of 10 µM caffeine (mean ± SD, n=3).

**(G and H)** Homodimerization of acVHH-V106D (G) and acVHH-WT (H). Binding of soluble acVHH-V106D or acVHH-WT (i.e. the binders) to acVHH variants displayed on yeast cells was assessed in the presence or absence of 10 µM caffeine (mean, n=2).

**(I)** Intracellular binding in the absence or presence of 30 µM caffeine was measured with the NanoBiT® protein complementation assay in Jurkat T cells (mean ± SD, n=3). These data are also part of Figure 1J.

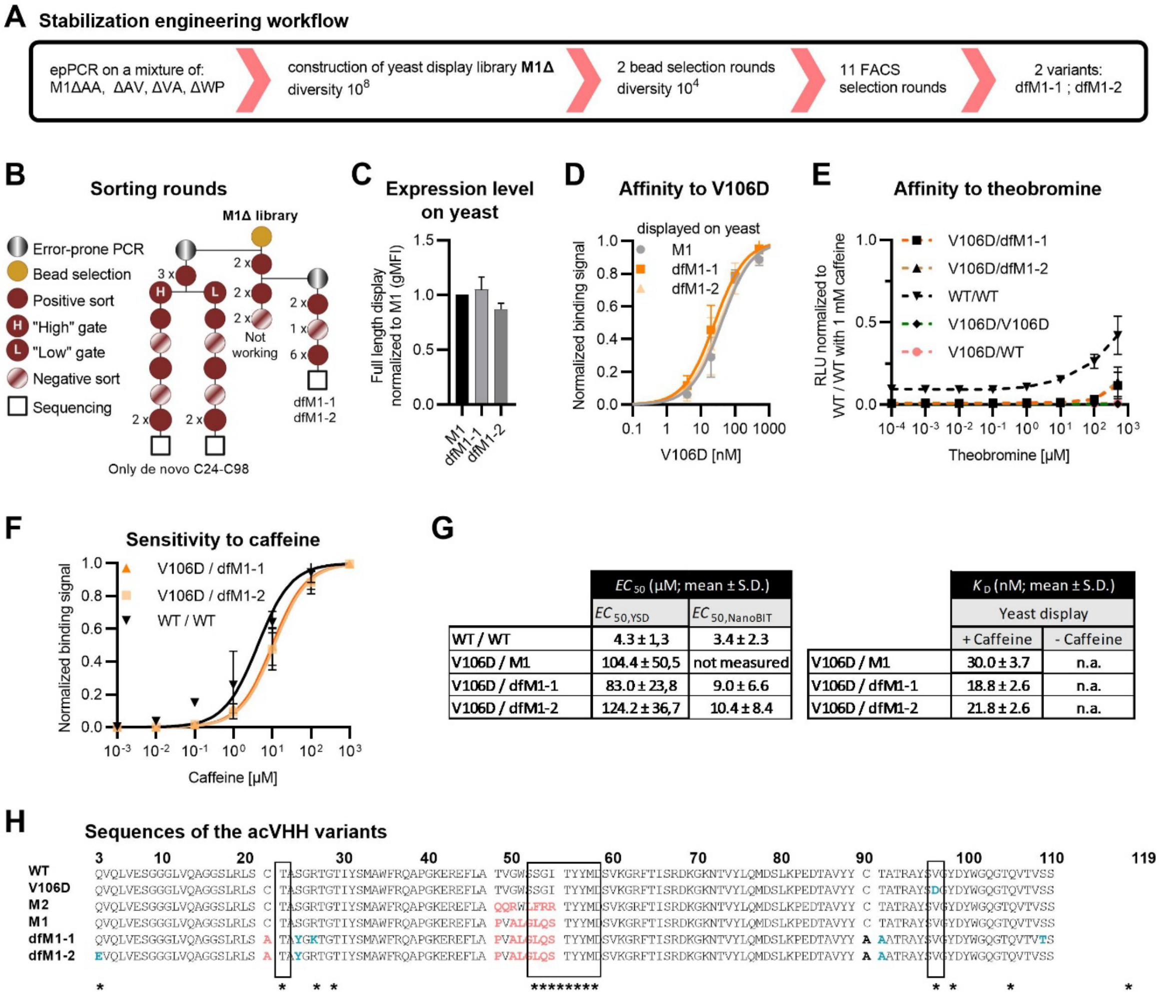


| | $EC_{50}$ (µM; mean ± S.D.) | |
|---|---|---|
| | $EC_{50,YSD}$ | $EC_{50,NanoBIT}$ |
| WT / WT | 4.3 ± 1,3 | 3.4 ± 2.3 |
| V106D / M1 | 104.4 ± 50,5 | not measured |
| V106D / dfM1-1 | 83.0 ± 23,8 | 9.0 ± 6.6 |
| V106D / dfM1-2 | 124.2 ± 36,7 | 10.4 ± 8.4 |

| | $K_D$ (nM; mean ± S.D.) | |
|---|---|---|
| | Yeast display | |
| | + Caffeine | - Caffeine |
| V106D / M1 | 30.0 ± 3.7 | n.a. |
| V106D / dfM1-1 | 18.8 ± 2.6 | n.a. |
| V106D / dfM1-2 | 21.8 ± 2.6 | n.a. |

**Supplemental Figure 2: Engineering of disulfide-free acVHH-variants.**

**(A)** Schematic of the stabilization engineering workflow. An equimolar mixture of M1ΔAA, ΔAV, ΔVA and ΔWP was subjected to error-prone PCR to create the M1Δ library, followed by 2 magnetic bead selections and 11 FACS selection rounds.

**(B)** Detailed schematic of the selection strategy. Additional rounds of error-prone PCR were performed as indicated to increase library diversity.

**(C)** Full-length display levels on yeast cells of disulfide-free acVHH-M1 variants. The gMFI was normalized to acVHH-M1 (mean ± SD, n=3).

**(D)** Binding of yeast-displayed acVHH mutants to increasing concentrations of soluble acVHH-V106D was assessed in the presence of 30 µM caffeine. The data for dfM1-1 are also depicted in Figure 1G.

**(E)** Intracellular binding of the acVHH variant dimers in the presence of increasing concentrations of theobromine was measured with the NanoBiT® protein complementation assay in Jurkat T cells. The signal (RLU) was normalized to the acVHH-WT homodimer in the presence of 1 mM caffeine (mean ± SD, n=3).

**(F)** Sensitivity to caffeine measured in the intracellular protein complementation assay shown in Figure 1K (mean ± SD, n=4). Data from Fig. 1K were background-subtracted and normalized to maximum binding signal to yield these dose response curves.

**(G)** Summary of $EC_{50}$ (sensitivity to caffeine) and $K_D$ values (affinities between the two nanobodies in the presence of caffeine) for all variants and different types of assays. Non analyzable data (n.a.).

**(H)** Sequence alignment of the different acVHH variants. Boxes and red indicate engineering spots, either the cysteines or the 8 amino acids (8NNK) loop. Point mutations outside of these positions are indicated in blue.

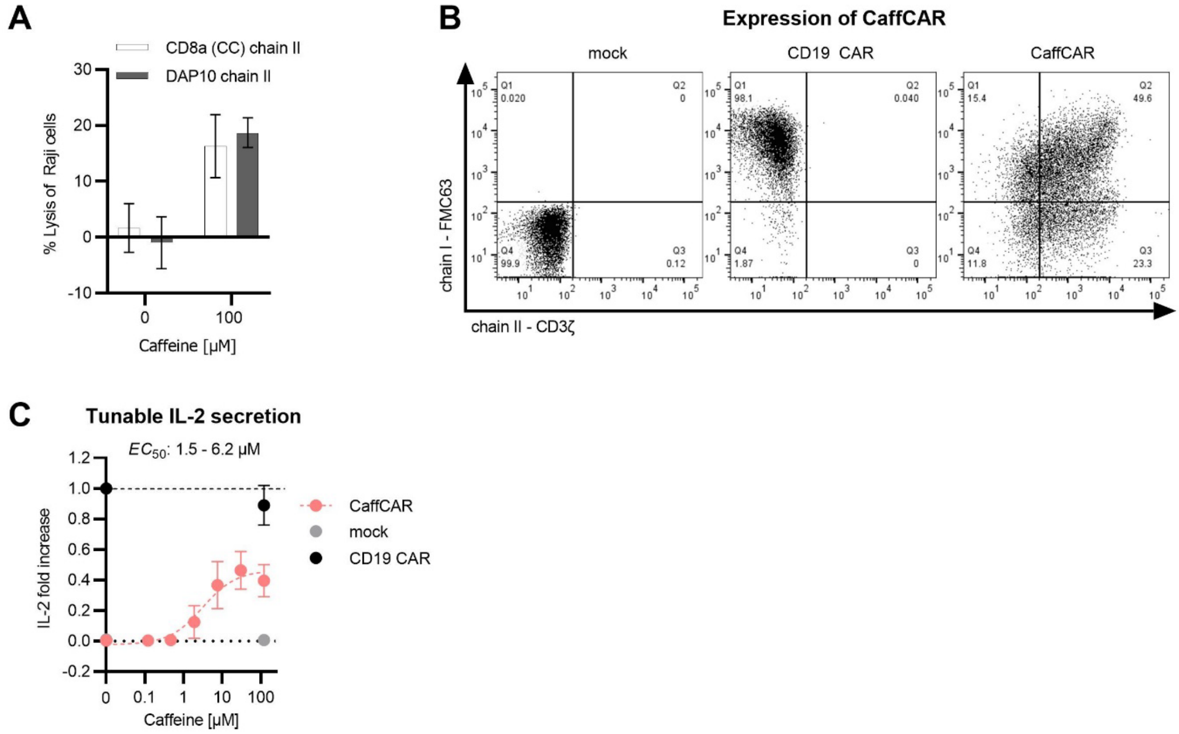


**Supplemental Figure 3: Split CAR regulation with caffeine**.

**(A)** Comparison of the cytotoxicity of CAR T cells containing different dimeric chain II variants. In both cases the chain I was monomeric and contained the dfM1-1 nanobody. The chain II either contained the DAP10 domain as depicted in Figure 2A or the CD8α hinge with cysteines as in Figure 2B. The lysis of Raji cells (E:T of 2:1, 24 h) was normalized to mock T cells without caffeine (n = 3).

**(B)** Expression of the CaffCAR compared to mock T cells and CD19 CAR.

**(C)** IL-2 secretion of CaffCAR T cells depending on caffeine concentration. Co-cultures were set up with an E:T ratio of 2:1 for 24h. IL-2 secretion was normalized to that of CD19 CAR without caffeine (n = 5, 5 different T cell donors). The data points for 0 and 120 µM caffeine were also used in Figure 2B.

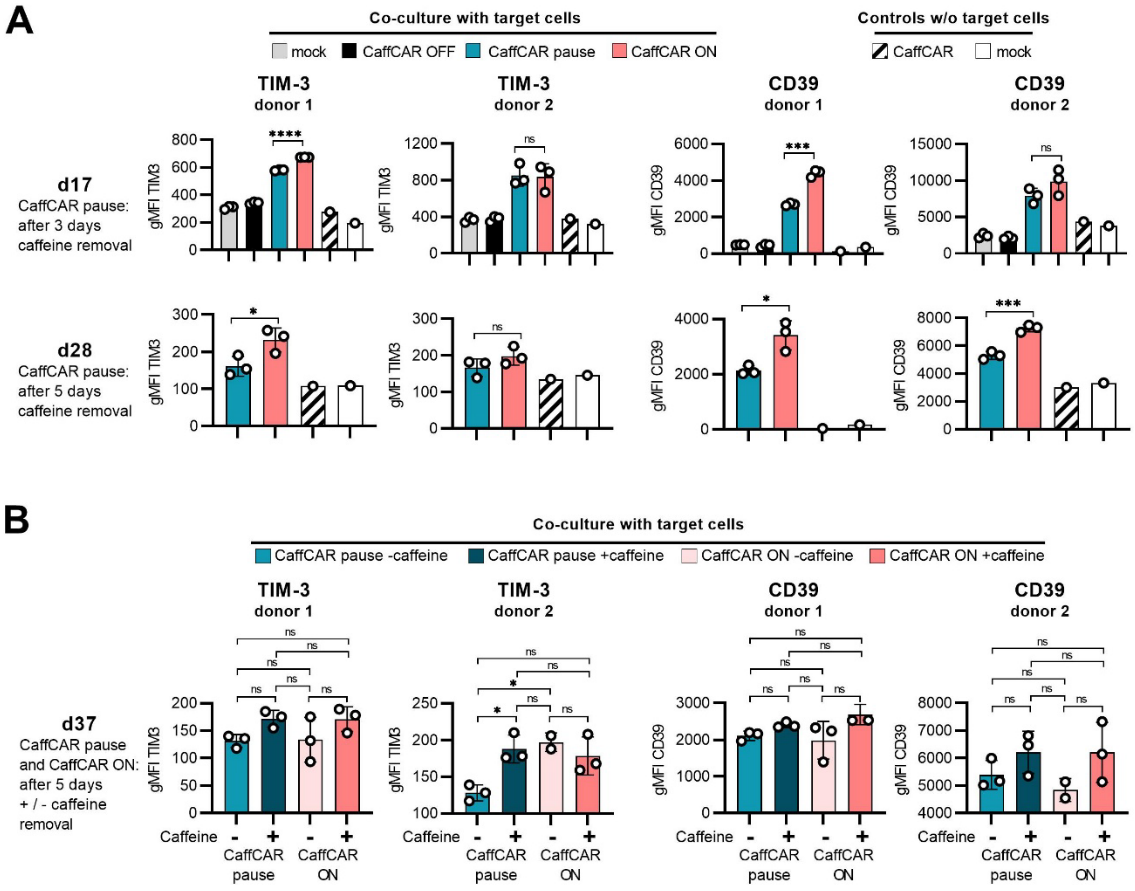


**Supplemental Figure 4: Pausing caffeine administration limits T cell exhaustion.**

**(A-B)** Expression of TIM-3 and CD39 on d17 and d28 **(A)** and d37 **(B)** of the long-term co-culture with Raji target cells. Technical triplicates for two independent T cell donors are shown.

* < 0.05; ** < 0.01; *** < 0.001; **** < 0.0001. Multiple unpaired t-test with Welsh's and Holm Sidak correction (A), and 1-way Anova with Turkey's test for d37 (B).

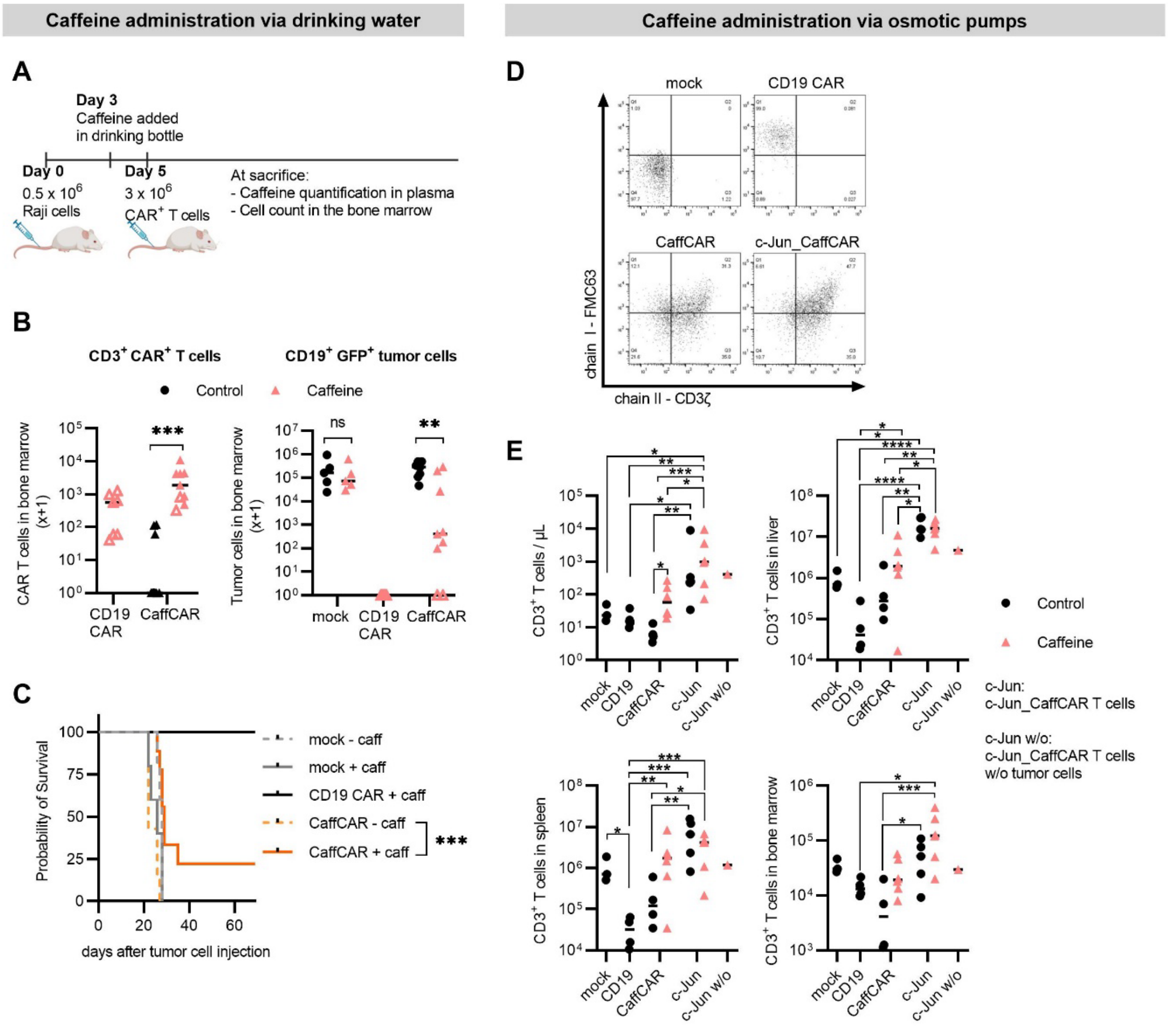


**Supplemental Figure 5: Caffeine regulated CaffCAR function *in vivo*.**

**(A-C)** *In vivo* experiment with caffeine administration via drinking water.

**(A)** Schematic of the experiment.

**(B**) Quantification of CAR T cells (left) and tumor cells (right) in the bone marrow of mice at sacrifice. The empty triangles represent mice sacrificed at the end point.

**(C)** Survival curves.

**(D-E)** *In vivo* experiment with caffeine administration via osmotic pumps.

**(D)** CAR expression of CaffCAR, c-Jun_CaffCAR and CD19 CAR T cells.

**(E)** Quantification of total T cells in circulation on d18 and in the liver, spleen, bone marrow at sacrifice (d22).

* < 0.05; ** < 0.01; *** < 0.001; **** < 0.0001. Multiple unpaired t test with Welch correction and Holm-Sidak corrected (B and E), log-rank Mantel-Cox test (C) and 2-way Anova with Turkey'test (F)
